# Agroforestry reshapes soil bacterial communities to enhance *Ricinus communis* oil quality and bioactivity over monocropping: comparative metagenomics and culture-dependent insights

**DOI:** 10.64898/2026.04.24.720747

**Authors:** Krishna Saharan, Pradeep Pagaria, Taskeen Bano, Nishant Chauhan, Rabisankar Mandi, Anas Mohd, Ravi Jameriya, Ramesh Kumar, Praduman Yadav, Pooja Yadav, Naincy Rathee, Narsingh Chauhan, Ayan Sadhukhan

## Abstract

An agroforestry (AF) system improves crop quality, ecosystem services, and microbial resilience, but its effects on oilseed bioactivity and soil microbiomes are still underexplored. This study compared AF and monocropping systems for castor (*Ricinus communis* L.) grown in Rajasthan, India, to evaluate plant productivity, seed oil composition, antimicrobial properties, and soil bacterial communities. AF enhanced seed morphology and germination. Castor oil from agroforestry had a 2.4-fold higher phenolic content, 2% more ricinoleic acid, and lower levels of oleic and linoleic acids compared to monocropping, confirmed by infrared spectroscopy and gas chromatography, along with increased expression of the *RcDGAT2* gene involved in fatty acid biosynthesis. This led to improved antimicrobial activity against *Bacillus mobilis* and *Pseudomonas fluorescens*. Full-length 16S rRNA gene sequencing on the Nanopore platform identified 17 bacterial phyla in soil microbiomes, with Proteobacteria and Firmicutes as the dominant phyla. While alpha diversity was similar, AF soils showed distinct taxonomic shifts, enriching bacteria such as *Alkalimonas*, *Aureimonas*, *Blastopirellula*, *Glutamicibacter*, *Rhizobium*, *Rhizomicrobium*, and *Rhodovulum*, linked to nutrient cycling and plant growth promotion. Isolated rhizospheric/root endophytic *Bacillus safensis* and *Enterobacter cloacae* from AF castor exhibited plant growth-promoting traits via biochemical tests and whole-genome sequencing; their oil biosynthesis genes likely contribute to host oil quality by enhancing precursor supply and phenolic pathways. These isolates enhanced the growth of the model plant *Arabidopsis thaliana*. In summary, AF enhances the bioactivity of castor oil and microbial functions by modulating plant-soil-microbe interactions, thereby supporting sustainable crop quality and soil health.

## 1. Introduction

Castor (*Ricinus communis* L.), a perennial shrub in the Euphorbiaceae family, holds significant agronomic and economic importance in India. Known as “Eranda” in Sanskrit, it is integrated into various farming systems, including in Rajasthan’s arid regions, which face water scarcity, high temperatures, and poor soils (Srinivasa Rao et al. 2012; Warra, 2015; Torrentes-Espinoza et al. 2017; Ramothloa et al. 2025). Native to Africa and India, castor thrives in tropical and subtropical areas, producing ricinoleic acid-rich seeds vital for biofuels, lubricants, pharmaceuticals, cosmetics, and antimicrobials, supporting livelihoods despite environmental challenges (Al-Mamun et al. 2016; Abomughaid et al. 2024). Versatile castor is grown in monocropping (MC) or agroforestry (AF) systems, covering 13.75 million hectares nationwide in states like Andhra Pradesh, Rajasthan, Maharashtra, and Uttar Pradesh. It is often paired with trees such as neem, babool, and acacia to improve soil health, biodiversity, climate resilience, and livelihoods (Sharma et al. 2017; Panwar et al. 2022). AF systems outperform MC by increasing productivity, conserving resources, promoting biodiversity, maintaining soil health, and stabilizing microclimates, while also helping reduce erosion, runoff, and the impacts of extreme weather in arid Rajasthan (Bulut and Gökalp, 2022; Panwar et al. 2022; Pancholi et al. 2023).

Cultivation systems affect growth, yield, seed traits, germination, and oil bioactives like ricinoleic acid and phenolics, conferring anti-inflammatory, antimicrobial, wound-healing, anti-HIV, anticancer, and antioxidant effects, with GC analysis revealing practice-driven variations (Torrentes-Espinoza et al. 2017; Kaur and Bhaskar, 2020; Mukhlis et al. 2022; Aneja et al. 2024). The environment shapes the soil microbiota, including endophytes that aid in growth, nutrient cycling, and disease resistance—where AF systems foster beneficial communities in degraded soils in Rajasthan (Tudi et al. 2021; Li et al. 2023).

Recent technological advancements, especially in metagenomics, have revolutionized the study of microbial communities. Full-length 16S rRNA gene sequencing, utilizing platforms such as Oxford Nanopore, enables the comprehensive profiling of soil microbiota at high resolution (Miranda-Carrazco et al. 2022). This method provides a comprehensive analysis of microbial communities, their functional capabilities, and ecological changes driven by different cropping systems. Understanding these microbial interactions is essential for advancing eco-friendly farming practices that leverage helpful bacteria to boost plant stress tolerance, minimize agrochemical use, and foster long-term soil health (Huang et al. 2022).

This study systematically evaluates the comparative impacts of AF and MC cultivation systems on castor under the arid agroclimatic conditions of Rajasthan, focusing on plant productivity, seed oil chemistry, antimicrobial functionality, and soil microbial ecology. The investigation integrates morpho-physiological assessments of seed traits and germination dynamics with detailed biochemical characterization of castor oil, including phenolic profiling and bioactivity evaluation against representative bacterial pathogens. To elucidate the responses of below-ground ecosystems, the soil microbial community structure and diversity are analyzed using full-length 16S rRNA gene sequencing, which enables high-resolution taxonomic and functional inference. By linking cultivation-induced shifts in plant biochemical traits with microbial community composition, this study aims to elucidate the mechanistic role of AF in enhancing plant–microbe interactions and ecosystem multifunctionality. The outcomes are expected to generate evidence-based insights that support AF as a resilient, resource-efficient, and ecologically sustainable production strategy for enhancing crop quality and enriching bioactive compounds in water-limited environments, such as Rajasthan.

## 2. Materials & Methods

### 2.1 Plant materials and sample collection

Castor (*Ricinus communis* L.) cultivar ICH-66, known for its high oil yield and favorable agronomic traits, was used in this study. These seeds were collected from two distinct agricultural cultivation systems: a demonstration plot maintained under an AF system and an MC system, both located at the Agriculture University, Jodhpur (AUJ), Rajasthan, India. AUJ has implemented an AF model on 3 acres of land at the University field site, with support from the Agroforestry Promotion Network (APN), Switzerland. The AF plot at AUJ features 20-year-old Rohida (*Tecomella undulata*) trees, while various crops, including fruit, timber, fodder, castor, and others, are grown in the interspaces. The plot is managed organically using indigenous cowdung-based fertilizers ‘Jeevamrit’ and ‘Beejamrit’ (Kaushal et al. 2024; Bhoi et al. 2025), along with dried Napier grass (*Cenchrus purpureus*) mulching, without the use of chemical fertilizers. During the selection process, healthy plants of similar age were randomly chosen to ensure they had well-developed primary and secondary branches, and the capsules were fresh and green. This was facilitated by marking the selected healthy plants with a red ribbon to consistently identify and differentiate them throughout the study.

### 2.2 Physical Properties of Castor

#### 2.2.1 Seed color and weight

The seed colors of both AF and MC-grown castor seeds were visualized and compared based on color darkness. For the 100-seed weight determination (Duaja et al. 2019), seeds were randomly selected, and at least four replicates of 100 pure seeds each were counted and weighed separately using an analytical balance.

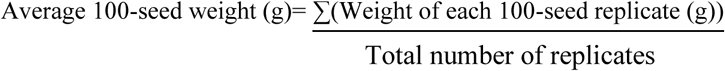

#### 2.2.2 Moisture content of Castor seeds

Twenty grams of pre-cleaned seeds were accurately weighed and oven-dried at 80°C for 6 h, with mass measurements taken at 2-h intervals. This process was continued until the weight stabilized. At each 2-h mark, samples were briefly removed from the oven, cooled in a desiccator for 20 min, and then re-weighed (Silva et al. 2020). Seed moisture percentage was determined using the following equation:

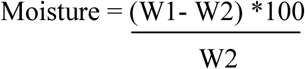

Here, W1 represents the initial mass of the sample prior to drying, while W2 denotes the mass after drying.

#### 2.2.3 Determination of castor seeds germination rate

Germination percentages for the AF and MC-grown seeds were determined via the paper towel assay (Pradhan and Lakpale, 2020). Seeds were positioned on wetted filter paper within petri dishes and checked daily. The average germination percentage was computed at the conclusion of the monitoring period. The germination index (GI) was computed using the formula: GI = ∑(nᵢ/tᵢ), in which nᵢ indicates the count of seeds germinated on day i, and tᵢ signifies the days post-sowing, with values summed across all days up to the cessation of germination.

### 2.3 Extraction and chemical analysis of castor oil

To extract the original consistency of castor oil, oil extraction was performed using a hydraulic press (Akhihiero et al. 2021). For each system, 100 kilograms of seeds were processed under standardized conditions to ensure comparability. The oil was extracted by hydraulic pressing, ensuring a cold-press process that preserves bioactive compounds. Physical parameters, including seed size, seed color, moisture content, and germination potential, were evaluated to determine morphological differences influenced by cultivation practices.

The chemical profile of the extracted castor oils was analyzed at the Indian Institute of Oilseeds Research in Hyderabad, India. Attenuated Total Reflectance-Fourier Transform Infrared spectroscopy (ATR-FTIR) was used to determine the composition and concentrations of key phenolic compounds and other bioactive constituents. This characterization provided essential baseline data, revealing how cultivation practices might influence the chemical makeup and potential bioactivity of the oils (Warra, 2015; Socaciu et al. 2020). Fatty acid methyl esters (FAMEs) were generated via a dual-step transesterification protocol as detailed by Yadav and Anjani (2017). In the initial phase, 100 mg oil samples were treated with 5 mL methanolic 2% H₂SO₄ at 50°C over 4 h. Post-reaction, the blend rested for 1 h to phase-separate, enabling removal of the overlying methanol-aqueous fraction. The bottom residue then received 2 mL 13% KOH in methanol and was incubated at 55°C for 30 min to complete base-catalyzed transesterification. Hexane-extracted organics were rinsed multiple times with water until pH neutrality, dried over anhydrous Na₂SO₄, and concentrated via N₂ purging to isolate FAMEs. Compositional analysis employed an Agilent 7890B GC coupled to FID and autosampler. Chromatography occurred on an HP-1 column (100% dimethylpolysiloxane; 30 m × 0.32 mm i.d., 0.25 μm film; Agilent). With N₂ carrier at 1.2 mL/min, the oven initiated at 150°C (2 min hold), ramped 10°C/min to 300°C. Samples (1 μL) injected at 100:1 split; inlet/detector at 325°C. Relative peak areas from Agilent EZChrom Elite Compact post-processing provided identification and % quantification.

### 2.4 Antibacterial activity testing of castor oil

To evaluate the antimicrobial activity of the castor oil, it was diluted in methanol to prepare a series of concentrations. The undiluted oil served as the original (100%), while dilutions at 10%, 30%, 50%, and 70% concentrations were prepared by mixing specific volumes of oil with methanol. For example, 100 μL of castor oil was mixed with 900 μL of methanol to achieve a 10% concentration, and similar adjustments were made for the higher concentrations (Alqahtani et al. 2019; Ibrahim and Kebede, 2020). These dilutions facilitated dose-response assessments in antimicrobial assays, as methanol acts as an effective solvent that ensures uniform dispersion of oil components in aqueous-based systems without exerting significant antimicrobial effects of its own. Ampicillin discs of 2 µg (Himedia) were used for the disc assay, while 50 µg mL^-1^ tetracycline (Sigma) was used for the well susceptibility assay and growth kinetics.

The bacterial strains selected for antimicrobial testing included the Gram-positive *Bacillus mobilis* and the Gram-negative *Pseudomonas fluorescens* (Paray et al. 2023), both isolated from the AF plot. These bacterial strains were isolated using serial dilution, and molecular identification was performed at the M.S. Swaminathan Research Foundation (MSSRF) in Chennai, India (Kannan et al. 2018; Al-Fadhal et al. 2019). These bacteria are representative of soil microbial communities and plant-associated bacteria, making them suitable models for assessing the antimicrobial potential of the castor oils within agricultural contexts.

The antimicrobial activity of the castor oils was tested using the agar well puncture method. In this assay, bacterial cultures grown overnight in Luria-Bertani broth were standardized to approximately 10^8^ CFU mL^-1^ and uniformly spread over sterilized LB agar plates. Using a sterile cork borer, four wells were punched into each agar plate, with two wells filled with 0.05 mL of each oil dilution, one well serving as a negative control (left empty), and another filled with tetracycline as a positive control (Govindan et al. 2022; Jena et al. 2025). Following incubation at 37°C for 24 h, the inhibition zones surrounding each well were quantified. Clear zones indicated the extent of bacterial growth suppression, providing a qualitative measure of antimicrobial efficacy. Additionally, the antibiotic assay involved impregnating sterile filter paper discs with various concentrations of castor oil, placing them on inoculated agar plates, and incubating them under similar conditions. The zones of inhibition around these discs were measured to compare the antimicrobial activity of castor oil with standard antibiotics (Hemeg et al. 2020).

For the growth kinetics assessment using the broth dilution method (Noval et al. 2019; Vanegas et al. 2021), six 100 mL conical flasks were labeled with oil concentrations of 10%, 30%, 50%, 70%, and 100% oil, along with an antibiotic as a control. Each flask received 50 mL of LB media, followed by the addition of 5 mL of the respective oil dilution and 100 µL of standardized bacterial inoculum. These flasks were incubated at 32°C on a shaker set to 120 rpm, with samples collected every 2 hours over 24 hours. Aliquots were transferred to sterile tubes for optical density measurements at 0, 2, 4, 6, 8, 10, and 24 hours. For the agar dilution method, phenol red (60 µL) was added to 90 mL of LB agar, and each oil series (10%, 30%, 50%, 70%, and 100%) was incorporated at a 9:1 ratio (90 mL agar to 10 mL castor oil). Petri dishes were each filled with about 15 mL of agar containing varying oil concentrations and left to solidify. Next, 50 µL of bacterial suspension was uniformly spread across the surfaces using a sterile swab, followed by incubation at 37°C for 24 h to evaluate growth inhibition. Post-incubation, plates were inspected for colony formation; the minimum oil level showing no visible growth was designated as the MIC (Mohammed et al. 2021).

### 2.5 Microbiome analysis of soil from castor rhizospheres

Soil samples were collected from the root zone of both MC and AF castor plants from the Agriculture University, Jodhpur (Kannan et al. 2018). Samples collected in polypropylene bags and kept in ice boxes before being carried to the laboratory (Sandilya et al. 2022). Soil DNA was isolated from samples using mechanical bead-beating, followed by purification with a commercial kit (DNeasy PowerSoil Pro, Qiagen). The 16S rRNA gene region was PCR-amplified using the 16S Barcoding Kit 24 V14 (SQK-16S114.24) according to the supplier’s guidelines. Amplicons were then cleaned up through a gel extraction procedure (GeneJET Gel Extraction Kit, Thermo Scientific). Libraries for sequencing were prepared with 16S Barcoding Kit 24 V14 (Oxford Nanopore Technologies, UK). Sequencing was performed on the FLO-MIN114 flow cell (Nanopore) using the default parameters. Sequence Base calling was performed using the high-quality option, with a minimum cutoff quality score of 9. Low-quality data (Q score < 9, read length < 800 bp and > 1500 bp) were discarded before phylogenetic analysis. 16S rRNA gene sequences from each sample (Table 1) were processed independently. The metagenomic data were analyzed using the Metagenomic Rapid Annotation using Subsystem Technology (MG-RAST) pipeline. Data analysis was performed using the EPIM2ME desktop suite with Kraken2 classifier for taxonomic assignment. The diversity and composition of soil microbiota were compared between MC and AF castor plants using various diversity indices and statistical tests, viz., analysis of variance (ANOVA) and *t*-test.

**Table 1.** Results of 16S rRNA gene sequencing from rhizospheric soil of castor from agroforestry and monocropping systems.

| Sample Description | Number of quality-filtered 16S rRNA gene sequences |  |  |  |  |  |
| --- | --- | --- | --- | --- | --- | --- |
| MC 1 | 6531 |  |  |  |  |  |
| MC 2 | 5544 |  |  |  |  |  |
| MC 3 | 4314 |  |  |  |  |  |
| AF 1 | 8523 |  |  |  |  |  |
| AF 2 | 6866 |  |  |  |  |  |
| AF 3 | 7216 |  |  |  |  |  |
| Apha diversity Indices | MC1 | MC2 | MC3 | AF1 | AF2 | AF3 |
| Berger Parker index | 0.09 | 0.11 | 0.06 | 0.05 | 0.07 | 0.06 |
| Effective number of species | 51.46 | 61.35 | 42.45 | 78.79 | 55.41 | 69.46 |
| Fisher's alpha | 16.9 | 17.72 | 10.39 | 19.68 | 14.39 | 17.89 |
| Inverse Simpson's index | 1.03 | 1.03 | 1.03 | 1.02 | 1.03 | 1.02 |
| Pielou's evenness | 0.86 | 0.9 | 0.94 | 0.93 | 0.92 | 0.93 |
| Richness | 96 | 96 | 55 | 107 | 79 | 96 |
| Shannon diversity index | 3.94 | 4.12 | 3.75 | 4.37 | 4.01 | 4.24 |
| Simpson's index | 0.97 | 0.97 | 0.97 | 0.98 | 0.98 | 0.98 |

MC: Monocropping field; AF: Agroforestry field; 1, 2, 3 are biological replicates

### 2.6 Isolation of rhizospheric and root endospheric bacteria from castor

Healthy plants of *R. communis* were selected for the collection of soil and root samples from AF and MC fields. Healthy root samples were randomly collected for isolation of root endophytes and transferred to zip-lock bags. These bags are labelled, sealed, brought to the laboratory, and stored at 4॰C in a deep freezer until isolation procedures are completed. One gram of rhizosphere soil from castor plants (AF and MC) at 30 cm depth was collected and processed using the standard serial dilution method. A rhizosphere soil sample was mixed with sterile phosphate-buffered saline (PBS) to create a series of tenfold dilutions (10⁻¹ to 10⁻⁹). A 100 μL volume of each dilution was then spread or poured onto Luria Bertani (LB) agar plates. The plates were incubated at 28°C for 24 h, and the number of colonies was recorded (CFU g^−1^). Morphologically distinct colonies were identified, subcultured, and maintained for further analysis.

### 2.7 Characterization of plant growth-promoting potential of bacterial isolates

Bacterial isolates were grown in peptone water broth for 48 hours at 30°C to assess ammonia production. Following incubation, 1 mL aliquots of each culture were transferred to microtubes and combined with 50 μL of Nessler’s reagent (prepared as 10% HgI₂, 7% KI, and 50% aqueous solution of 32% NaOH). A faint yellow color signaled low-level ammonia production, whereas a pronounced yellow-to-brownish coloration denoted high ammonia output (Dye et al. 1962). To evaluate indole acetic acid (IAA) production, isolates were cultured in LB medium supplemented with 500 mg L⁻¹ tryptophan at 30°C for 48 hours. Cultures underwent centrifugation, after which 1 mL supernatant was thoroughly blended with 2 mL Salkowski’s reagent and incubated in darkness at room temperature for 30 min. Development of a pink hue confirmed IAA synthesis (Sheng et al., 2008). For the zinc solubilization test the isolates were grown on the zinc solubilization agar (Himedia, M2068-500G) containing 10 g L^-1^ dextrose, 1 g L^-1^ (NH_4_)_2_SO_4_, 0.2 g L^-1^ KCl, 0.1 g L^-1^ K_2_HPO_4_, 0.2 g L^-1^ MgSO_4_, pH 7.0, 0.1 % insoluble zinc compounds (i.e., ZnCO_3_) and 15 g L^-1^ agar, at 30°C for 48 h. The presence of a clear halo surrounding the colonies signifies zinc solubilization (Saravanan et al. 2007). Phosphate solubilization ability of the bacterial isolates was evaluated by testing their capacity to dissolve inorganic phosphate. In the experiment, Ca_3_(PO4)_2_, the insoluble inorganic form of phosphate, was included in Pikovskaya agar medium prepared with the following components per liter: 0.5 g yeast extract, 5 g Ca_3_(PO₄)₂, 0.2 g (NH₄)₂SO₄, 1 g MgSO₄·7H₂O, 0.1 mg MnSO₄, 0.1 mg FeSO₄, 10 g glucose, and 20 g agar, with the pH adjusted to 7. After streaking onto plates, the bacteria were cultured for 5 days at 30°C. The bacterial colony’s ability to solubilize insoluble phosphate was demonstrated by the formation of a translucent halo around it (Pikovskaya et al. 1948). To evaluate the siderophore production ability of the bacterium, it was cultured on a medium supplemented with chrome azurol S (CAS) and hexadecyltrimethylammonium bromide (HDTMA). Siderophore secretion resulted in an orange halo around the colony, formed by the dye chelating iron (Schwyn and Neilands, 1987).

### 2.8 Whole genome sequencing of bacterial isolates

The bacterial isolates were grown in Luria–Bertani (LB) broth at 37 °C for 24 h under continuous shaking at 180 rpm min⁻¹. Genomic DNA was extracted from the cultured cells using the HiPurA® Bacterial Genomic DNA Purification Kit (Himedia, MB505-250PR) according to the manufacturer’s protocol, and its quality was evaluated by 0.8% (w/v) agarose gel electrophoresis. The genomic DNA of the four isolates was sequenced on the Oxford Nanopore platform, with sequencing libraries prepared using the Rapid Barcoding Kit 96 V14 (SQK-RBK114.96, Oxford Nanopore, UK) and run on the MinION Mk1B device according to the supplier’s instructions. Raw sequence data were base-called and quality-filtered (Q score > 9, barcode-trimmed) using the fast base-calling feature in MinKNOW (software ver. 25.09.16). The processed reads were then de novo assembled with Flye v2.9.6 (Kolmogorov et al. 2019) via the NanoGalaxy server (https://nanopore.usegalaxy.eu/) (de Koning et al. 2020), and the resulting draft genomes were evaluated using QUAST v5.3.0 (Gurevich et al. 2013). Genome annotation was performed using the RASTtk v1.073 pipeline (Brettin et al. 2015) on the KBase platform (https://www.kbase.us) (Wood-Charlson et al. 2026). Circular genome maps were generated using Proksee (Grant et al. 2023). Taxonomic assignment of the isolates was carried out using both 16S rRNA-gene-based phylogenetic analysis and whole-genome-based phylogenomics. The 16S rRNA gene sequences retrieved from the draft genomes were used to identify homologs in the NCBI nucleotide database (16S ribosomal RNA sequences, Bacteria and Archaea; Update date: 2026/04/08) via BLASTn. Genome-scale taxonomic classification was further refined using the Genome Taxonomy Database accessed through NanoGalaxy (https://nanopore.usegalaxy.eu/), yielding high-resolution taxonomic placement. Average nucleotide identity (ANI) values of the sequenced genomes were computed against phylogenetically related species in NCBI Genomes using FastANI v1.3 (Jain et al. 2018). Finally, functional categorization of genes was undertaken based on SEED Subsystems, focusing on genes associated with nutrient assimilation, colonization potential, lipid (oil)-producing pathways.

### 2.9 Plant growth-promotion assay

Overnight-grown bacterial cultures were centrifuged at 5,000 rpm and resuspended in sterile PBS. *Arabidopsis thaliana* ecotype Col-0 seeds were surface sterilized by ethanol and sodium hypochlorite and imbibed in sterile PBS (control) or in PBS containing bacteria for 3 days at 4°C. Then the seeds were sown on nylon meshes (pore size N50), which were floated on ¼-Hoagland’s solution using photographic mounts and Styrofoam. The seeds were allowed to germinate and grow for 14 days, then transferred to a sterilized 1:2:1 mixture of soil, soil rite, and vermiculite in 2-inch pots (Verma et al. 2025). The treatment pots were inoculated with 1 mL of bacterial culture resuspended in PBS, four times at 5-day intervals, while the control boxes were mock-inoculated with 1 mL sterile PBS. Plants were imaged following one month of cultivation in a growth chamber set to 22°C, 60% relative humidity, 40 μmol m⁻² s⁻¹ light, and a 12/12 h light/dark photoperiod.

### 2.10 Real-time PCR analysis of gene expression

Developing spiny fruits (regmata) of castor were collected at three stages from AF and MC plots and flash-frozen in liquid nitrogen. They were crushed into a fine powder with a mortar and pestle. RNA was isolated using a CTAB-containing extraction buffer with the addition of 4% w/v PVP and 10% v/v β-mercaptoethanol. Following extraction, the samples were loaded onto the SV Total RNA Isolation System (Promega) columns for further purification. Conversion to cDNA was performed using RevertAid reverse transcriptase and random hexamer primers (Thermo Scientific). To assess expression levels of castor fatty acid biosynthesis genes, RT-qPCR was performed using the synthesized cDNA as template, TB Green Premix Ex Taq II (Tli RNase H Plus) (Takara), and primers detailed in Table S1. Three biological replicates were used for each sample, and *RcUBQ* was used as the housekeeping gene for normalization. The primer sequences were obtained from Rodríguez-Cabal et al. (2020).

### 2.11 Statistical Analysis

The data collected from all experiments were subjected to rigorous statistical analysis using SPSS Version 27 on a Windows 10 platform. A one-way ANOVA followed by post-hoc Tukey’s honestly significant difference (HSD) test was used to detect significant differences among the treatment groups at *P* < 0.05.

## 3. Results

### 3.1 Seed characteristics of castor from agroforestry and monocropping systems

The physical evaluation of the seeds revealed notable differences between the two cultivation systems. Seeds from the AF system were observed to be larger, healthier, darker, and more robust, while seeds from the MC system appeared smaller, lighter, and less vigorous. The yield from the agroforestry seeds was approximately 35 L of castor oil, while the MC seeds yielded about 32 L, reflecting differences in seed quality and oil content. In terms of weight variability, seeds from AF systems are heavier than those from MC systems. Higher moisture content is found in seeds from MC systems compared to those from AF systems. The average moisture content in the AF system is 63.33%, which is slightly lower than the 65.10% observed in the MC system. The average germination rate in the AF system is 69%, whereas in the MC system it is 66%. The Germination Index (GI) is 2.47 for the AF system and 2.45 for the MC system (Fig. 2A).

**Fig. 1.**
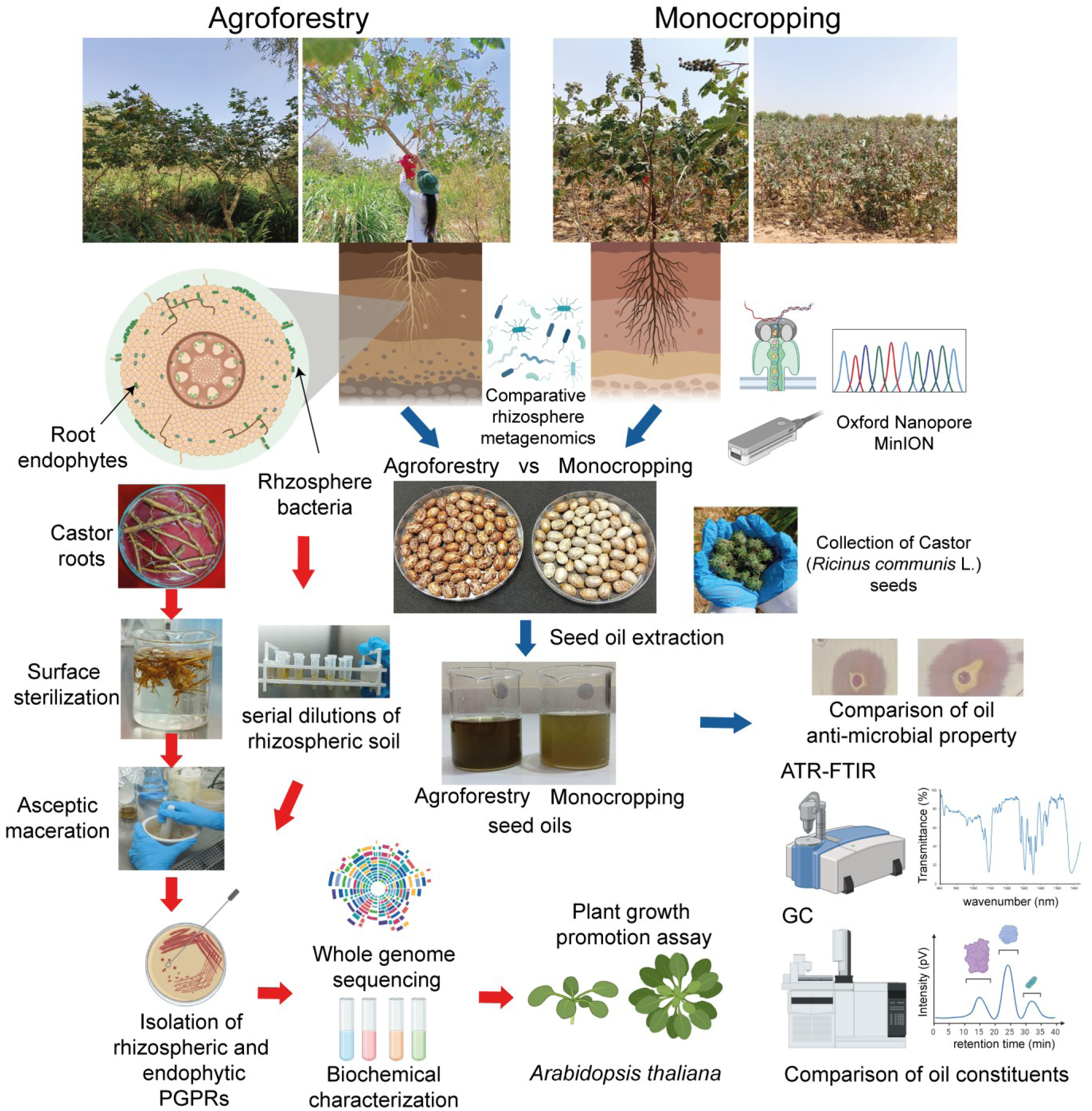
Schematic diagram of the comparative analysis of agroforestry versus monocropping systems for castor. Seeds of castor (*Ricinus communis* L.) plants, grown in agroforestry and monocropping systems at Agriculture University, Mandor, Rajasthan, were collected and compared for seed oil characteristics using attenuated total reflectance-Fourier transform infrared spectroscopy (ATR-FTIR) and gas chromatography (GC), and antimicrobial activity against *Bacillus mobilis* and *Pseudomonas fluorescens*. Parallel to this, roots from agroforestry-grown castor plants were collected, surface-sterilized, and crushed to isolate endophytic bacteria. Serial dilutions of soil around the roots were also used to isolate rhizospheric bacteria. The pure bacterial isolates were characterized using biochemical tests, whole-genome sequencing, and growth assays in *Arabidopsis thaliana* to assess their plant growth-promoting properties. Figure created with BioRender (https://www.biorender.com/).

**Fig. 2.**
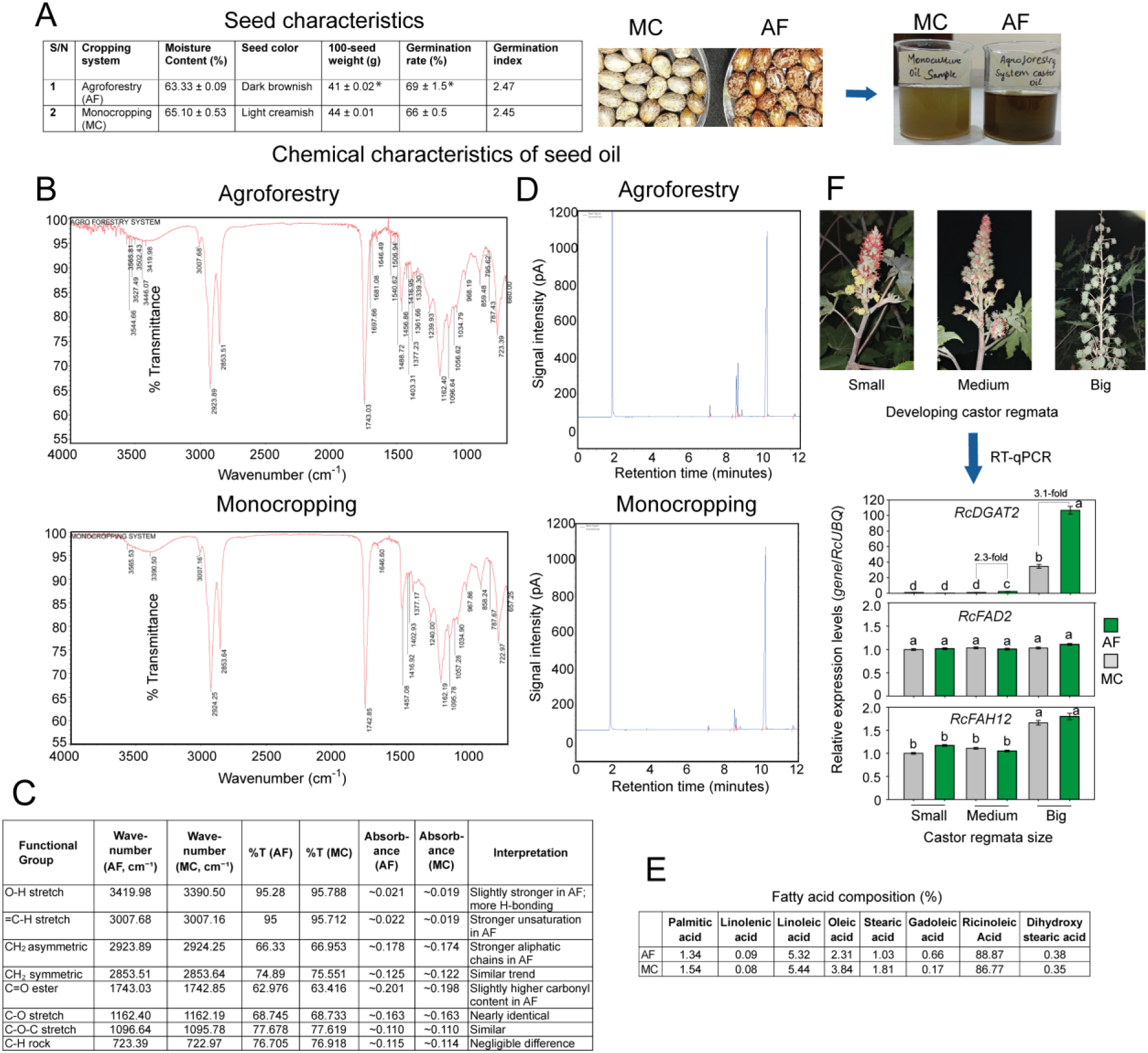
Physicochemical characteristics of castor grown in agroforestry versus monocropping systems. (**A**) The physical properties and germination rates of castor seeds are presented for agroforestry (AF) and monocropping (MC) systems. The data shown are the average of four replicates with standard error. An asterisk indicates a significant difference between AF and MC. (**B**) The Attenuated total reflectance-Fourier transform infrared spectroscopy (ATR-FTIR) spectra are shown as plots of % transmittance versus wavenumbers (cm^-1^). (**C**) Details of the functional groups in seed oil AF and MC systems were analyzed through infrared spectroscopy. (**D**) Gas chromatography (GC) spectra are shown as plots of signal intensity (pV) versus retention time in minutes. (**E**) The fatty acid compositions (%) of castor oil from AF and MC systems, analyzed through GC, are shown (average of three replicates). (**F**) Relative expression analysis of fatty acid biosynthesis pathway genes (*RcDGAT2*, *RcFAD2*, and *RcFAH12*) in castor regmata of three different stages from MC and AF plots. *RcUBQ* was used as a housekeeping gene for normalization. The data shown are the average of three biological replicates with standard errors. Different letters above the bars indicate significant differences (One-way ANOVA followed by Tukey’s test, *p* < 0.05).

### 3.2 Castor oil from agroforestry shows differences in chemical composition from monocropping

ATR-FTIR spectra of castor oil from MC and AF system displayed the characteristic signatures of ricinolate triglycerides, including ν(C–H) symmetric at 2923.89/2924.25 and asymmetric at 2853.51/2853.64cm⁻¹, ν(C=O) ester at 1743.03/1742.85 cm⁻¹, δ(CH₂/CH₃) at 1465/1377 cm⁻¹, a broad ν(O–H) envelope at 3419.98/3390.50 cm⁻¹, and C–O stretching bands at 1162.40/1162.19 cm⁻¹. Relative to MC, AF exhibited higher intensities in the aromatic/phenolic windows (1605–1505 and 1240–1220 cm⁻¹), together with a slightly more pronounced O–H shoulder, consistent with an increased contribution of phenolic/aromatic metabolites. Conversely, MC showed a relatively stronger 1065–1030 cm⁻¹ band attributable to the secondary-alcohol C–O of ricinoleate. These differences suggest that the AF system yields castor oil with a higher phenolic load and potentially greater antioxidant capacity, while the MC oil expresses a comparatively stronger ricinoleate alcohol signature (Fig. 2B, C). The ATR-FTIR instrument was pre-calibrated for the target compounds using partial least squares (PLS) and principal component analysis (PCA) with authentic standards. Phenolic compounds were identified by peak-ratio analysis and quantified by measuring peak area at the specified retention time. This method indicated that castor oil from AF had a higher phenolic content of 44 mg gallic acid equivalent (GAE) g⁻¹ than that from MC (18 mg GAE g⁻¹).

GC analysis of castor oil from AF and MC also showed compositional differences. Both chromatograms showed similar fatty acid elution patterns (Fig. 2D). A strong solvent peak was observed at ∼0.5 min. A major peak at 8.5–9 min corresponded to ricinoleic acid, the hallmark component of castor oil, while minor fatty acid peaks (palmitic, stearic, oleic, and linoleic acids) were observed between 5–8 min. Castor oil from AF contained 2% higher ricinoleic acid, 0.01% higher linolenic acid, and 0.49% higher gadoleic acid (Fig. 2E). On the other hand, oil from MC contained 0.12% higher linoleic acid, 0.78% stearic acid, 1.53% oleic acid, and 0.2% higher palmitic acid. The higher ricinoleic acid and lower oleic and linoleic acid contents in the AF sample suggest superior oil quality for industrial applications. To investigate the genetic basis of differences in oil properties between AF and MC, we estimated the relative expression levels of three fatty acid biosynthesis genes *diacylglycerol acyltransferase* (*RcDGAT2*), *oleoyl-12-desaturase* (*RcFAD2*), and *Oleoyl-12-Hydroxylase* (*RcFAH12*) (Héctor et al. 2020) in the total RNA extracted from developing castor regmata of different stages. The results indicated that while *RcFAD2* and *RcFAH12* levels were similar between AF and MC samples, *RcDGAT2* showed significantly higher expression in medium (2.3-fold) and big (3.1-fold) AF-grown castor regmata than in those from MC (Fig. 2F).

### 3.3 Castor oil from agroforestry shows greater antimicrobial potential than monocropping

#### 3.3.1 Disc susceptibility assay

The antimicrobial activity of castor oil against *Bacillus mobilis* and *Pseudomonas fluorescens* was assessed through zones of inhibition (ZOI) at varying concentrations, revealing that oils derived from AF systems exhibited significantly greater efficacy than those from MC systems (Fig. 3A). For *Bacillus mobilis*, the highest ZOI was observed at a 10% oil concentration of AF-derived castor oil (7.20 mm), with a gradual decline at higher concentrations. In contrast, MC-derived oil showed considerably lower activity, with ZOIs ranging from 0.83 mm (100% oil) to 1.25 mm at 70%. Similarly, *Pseudomonas fluorescens* was more susceptible to AF oil, with maximum inhibition at 10% concentration (5.53 mm) and 30% (5.77 mm), and reduced zones at higher concentrations. In contrast, MC oil demonstrated minimal activity across all concentrations.

**Fig. 3.**
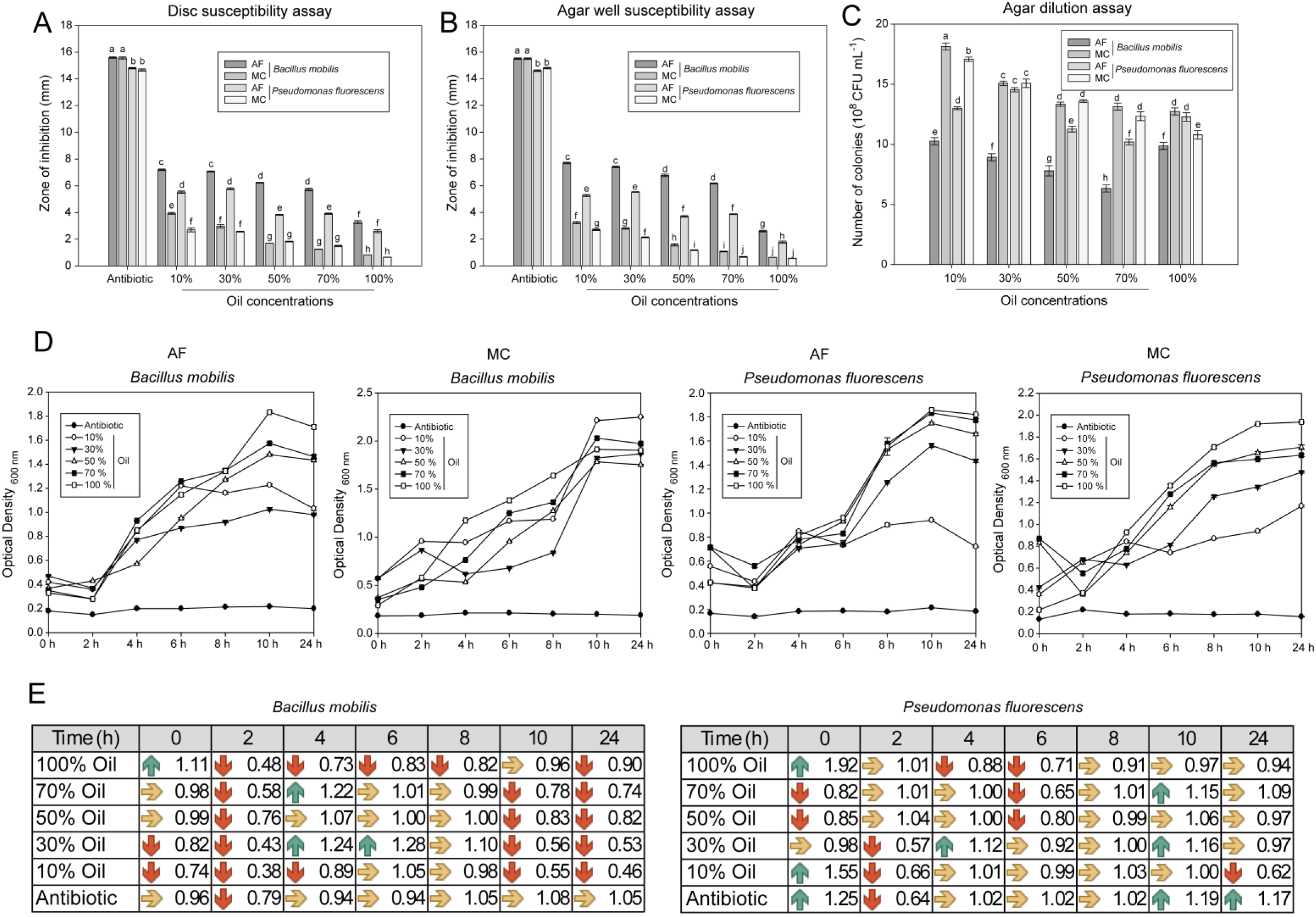
Comparison of the antibacterial properties of castor oil from agroforestry and monocropping systems. The antibacterial activities of different concentrations of seed oil (10%, 30%, 50%, 70%, and 100%) from agroforestry (AF) and monocropping (MC) systems against *Bacillus mobilis* and *Pseudomonas fluorescens* in various assay formats are presented. (**A**) Disc susceptibility assay. (**B**) Agar well susceptibility assay. (**C**) Agar dilution assay. Different letters above the bars indicate significant differences (One-way analysis of variance followed by post-hoc Tukey’s HSD test; *P* < 0.05; *N* = 3 replicates). (**D**) Growth curves of *Bacillus mobilis* and *Pseudomonas fluorescens* at various time points post-inoculation in Luria-Bertani broth are shown as optical density at 600 nm. The data shown are the average of three replicates with standard error. The Ampicillin discs (2 𝜇g) were used as controls for the disc susceptibility assay, while tetracycline (50 𝜇g/mL) was used for the agar well susceptibility, agar dilution, and growth kinetics assays. (**E**) The table shows the change in growth kinetics due to treatment with castor seed oil from AF, relative to that from MC. A value greater than 1.1 indicates faster growth (higher optical density) of the respective AF treatment, indicated by green "up-arrow"; a value below 0.9 indicates slower growth (lower optical density), indicated by red "down-arrow", and values between 0.9 and 1.1 are considered comparable, indicated by a yellow "right-arrow".

#### 3.3.2 Agar well susceptibility assay

The data from the agar well susceptibility assay indicate that castor oil derived from AF systems exhibits stronger antimicrobial activity compared to oil from MC systems against both *Bacillus mobilis* and *Pseudomonas fluorescens* (Fig. 3B). Specifically, *Bacillus mobilis* treated with AF oil showed larger zones of inhibition, ranging from 2.60 mm at a 100% concentration to a maximum of 7.70 mm at a 10% concentration, with a slight decrease at higher concentrations. Similarly, *Pseudomonas fluorescens* exhibited larger inhibition zones with AF oil, peaking at 5.53 mm at 30% concentration. Conversely, MC-derived oils consistently demonstrated lower zones of inhibition across all concentrations, with *B. mobilis* zones ranging from 0.63 mm (100%) to 3.23 mm (10%), and *P. fluorescens* (MC) zones from 0.56 mm (100%), 2.70 mm (10%), to 0.67 mm (070%). The antibiotic control produced significantly larger zones (approximately 14.6–15.5 mm), confirming the assay’s validity.

#### 3.3.3 Agar dilution assay

The agar dilution assay results demonstrate that bacterial growth, measured in 10^8^ colony-forming units (CFU)/mL, generally decreases as castor oil concentration increases across all tested groups (Fig. 3C). In the AF *B. mobilis* group, the bacterial count was highest at the 10% concentration (10.27 ± 0.29) and decreased progressively to its lowest at the 70% concentration (6.33 ± 0.29), before slightly increasing again at 100% (9.87 ± 0.29). Similarly, the AF *P. fluorescens* group showed a reduction from 13.00 ± 0.12 at 10% to 10.20 ± 0.23 at 70%, with a slight rise at 100%. The MC *B. mobilis* group started with a high bacterial count at 10% (18.13 ± 0.29) and declined at higher concentrations, with counts of 15.07 ± 0.18 at 30%, 13.33 ± 0.18 at 50%, and 13.13 ± 0.29 at 70%, before decreasing further at 100%. The MC *P. fluorescens* group exhibited a similar pattern, with counts decreasing from 17.07 ± 0.18 at 10% to 12.33 ± 0.37 at 70%. Overall, the data indicate a concentration-dependent inhibitory effect of castor oil on bacterial growth, with the most significant reductions observed at intermediate concentrations (50% and 70%) in the AF groups. The MC groups tend to maintain higher bacterial counts across all concentrations, suggesting slightly lower susceptibility to the antibacterial effects of MC-derived castor oil.

#### 3.3.4 Growth kinetics assessment

Higher concentrations of castor oil tend to increase the growth rate (Optical density, OD) in all bacterial systems, indicating enhanced bacterial growth or activity (Fig. 3D). For *Bacillus mobilis* and the AF-derived oils, the 70% concentration shows the most significant increase in optical density, with a value of 1.35 after 8 hours and 1.57 peaking around 10 hours. In contrast, the 10% and 30% concentrations inhibit bacterial growth most significantly at the same time point, with values of 1.17 and 0.92, respectively. Similarly, in *Pseudomonas fluorescens*, across the same AF oils, the 70% and original concentrations yield higher optical densities (1.83 and 1.86) and less inhibition than lower concentrations, with notable growth observed at 8 hours and beyond. In the MC systems, both *Bacillus mobilis* and *Pseudomonas fluorescens* show a progressive increase in optical density with increasing castor oil concentration, reaching their maximum levels around 10–12 hours. The 70% and original concentration again demonstrate the most pronounced growth, 1.60 and 1.92 in MO PF, while 2.03 and 1.91 in MO BM, suggesting that this concentration promotes bacterial activity more effectively than lower doses, while 10%, 30% and 50% show inhibition more frequently. Conversely, the antibiotic treatment maintains relatively low optical densities across all systems, indicating inhibition of bacterial growth. Fig. 3E compares the growth kinetics data for *B. mobilis* and *P. fluorescens* treated with castor oil from the AF plants with that from the MC plants. A value greater 1.0 indicates faster growth (higher OD) of the respective AF field treatment, indicated by green "up-arrow" (threshold: 1.1), a value below 1.0 indicates slower growth (lower optical density), indicated by red "down-arrow" (threshold: 0.9) and values between 0.9 and 1.1 are considered as similar, indicated by a yellow "right-arrow". The data for *B. mobilis* show that in 50% of cases, slower growth (down arrows) occurs; in 40% of cases, similar growth (right arrows) is observed; and in 10% of cases, faster growth is observed in treatments with oil from AF. For *P. fluorescens*, the data indicate that in 24% of cases, slower growth (down arrows) occurred; in 57% of cases, similar growth (right arrows) was observed; and in 19% of cases, faster growth was observed in treatments with oil from AF.

### 3.4 16S rRNA gene sequencing reveals unique bacterial taxa associated with castor in agroforestry and monocropping systems

Full-length 16S rRNA gene sequences were successfully amplified using the Nanopore 16S Barcoding Kit and subjected to quality filtering. The number of high-quality sequences per sample ranged from 4,314 to 8,523 (Table 1). Diversity indices across all samples revealed considerable microbial richness, with no statistically significant differences detected between the test and control groups (Table 1). The alpha diversity indices, including Shannon diversity, Simpson’s index, and richness measures, demonstrated similar community complexity across samples. Sequences were classified into 17 bacterial phyla. The dominant phyla included *Proteobacteria*, *Firmicutes*, *Actinobacteria*, *Planctomycota*, *Acidobacteria*, and *Bacteroidota*, which together accounted for over 60% of the community (Figure 4A). A minor fraction (<1%) belonged to *Verrucomicrobia*, *Armatimonadetes*, *Deinococcus-Thermus*, and other phyla (Figure 4B). Although some variation in relative abundance was noted between groups, these differences were not statistically significant. The sequences were assigned to 55 bacterial classes (Figure 4C, D). The most abundant classes (>1%) included *Alphaproteobacteria*, *Bacilli*, *Planctomycetia*, *Deltaproteobacteria*, *Gammaproteobacteria*, *Betaproteobacteria*, and *Actinomycetia* (Figure 4C). Notably, higher relative abundances of *Alphaproteobacteria* and *Planctomycetia* were observed in the AF group, whereas *Bacilli*, *Rubrobacteria*, and *Clostridia* were more prevalent in MC, although differences were not statistically significant (Figure 4C, D). At the family level, 265 families were identified, with 20 families accounting for more than 1% of the sequences. Prominent families included *Bacillaceae*, *Methylobacteriaceae*, *Gemmataceae*, *Paenibacillaceae*, *and Rhizobiaceae.* Several families associated with nitrogen fixation and biogeochemical cycling, such as G*emmataceae, Nitrospiraceae, Hyphomicrobiaceae, Xanthomonadaceae, Nitrobacteraceae, Rhizobiaceae* and *Planctomycetaceae* showed higher abundance in the AF group, although statistically insignificant. Similarly, *Bacillaceae, Rubrobacteraceae*, and *Spirochaetaceae* were differentially abundant in the MC group, with only *Spirochaetaceae* showing statistical significance (Figure 4E). A total of 3,835 operational taxonomic units (OTUs) were identified across all soil samples, representing 710 bacterial genera. The ten most abundant genera—*Microvirga*, *Rubrobacter*, *Bacillus*, *Paenibacillus*, *Vicinamibacter*, *Priestia*, *Nitrospira*, *Domibacillus*, *Gemmata*, *Skermanella*, *Urbifossiella*, *Neobacillus*, and *Gemmatimonas*—collectively accounted for over 25% of the total bacterial diversity (Figure 4F). Further analysis revealed that 382 bacterial species were detected exclusively in one or two samples within the MC group. Among these, only three OTUs reached statistical significance, including *Cerasibacillus*, which is known to contribute to the degradation of organic matter and nutrient cycling under alkaline soil conditions (Table 2). Conversely, the AF group exhibited 39 unique OTUs that were significantly enriched in AF compared to MC. Notably, *Rhodovulum*, *Rhizomicrobium*, *Glutamicibacter uratoxydans*, *Blastopirellula*, *Dokdonella*, *Devosia humi, Limimonas halophila*, *Bythopirellula* were among the significant OTUs in AF samples, which are implicated in nitrogen cycling. Other significantly enriched taxa in AF included *Pseudenhygromyxa*, *Streptomyces liangshanensis*, *Rhizomicrobium electricum*, *Alkalimonas*, *Thermodesulfomicrobium*, and *Tranquillimonas rosea*, involved in sulfur cycling and organic matter decomposition, as well as *Glutamicibacter*, *Aureimonas*, *Paraburkholderia phenazinium*, and *Rhizobium cellulosilyticum*, involved in plant growth promotion (for details on statistically significant OTUs: refer to Table 2).

**Fig. 4.**
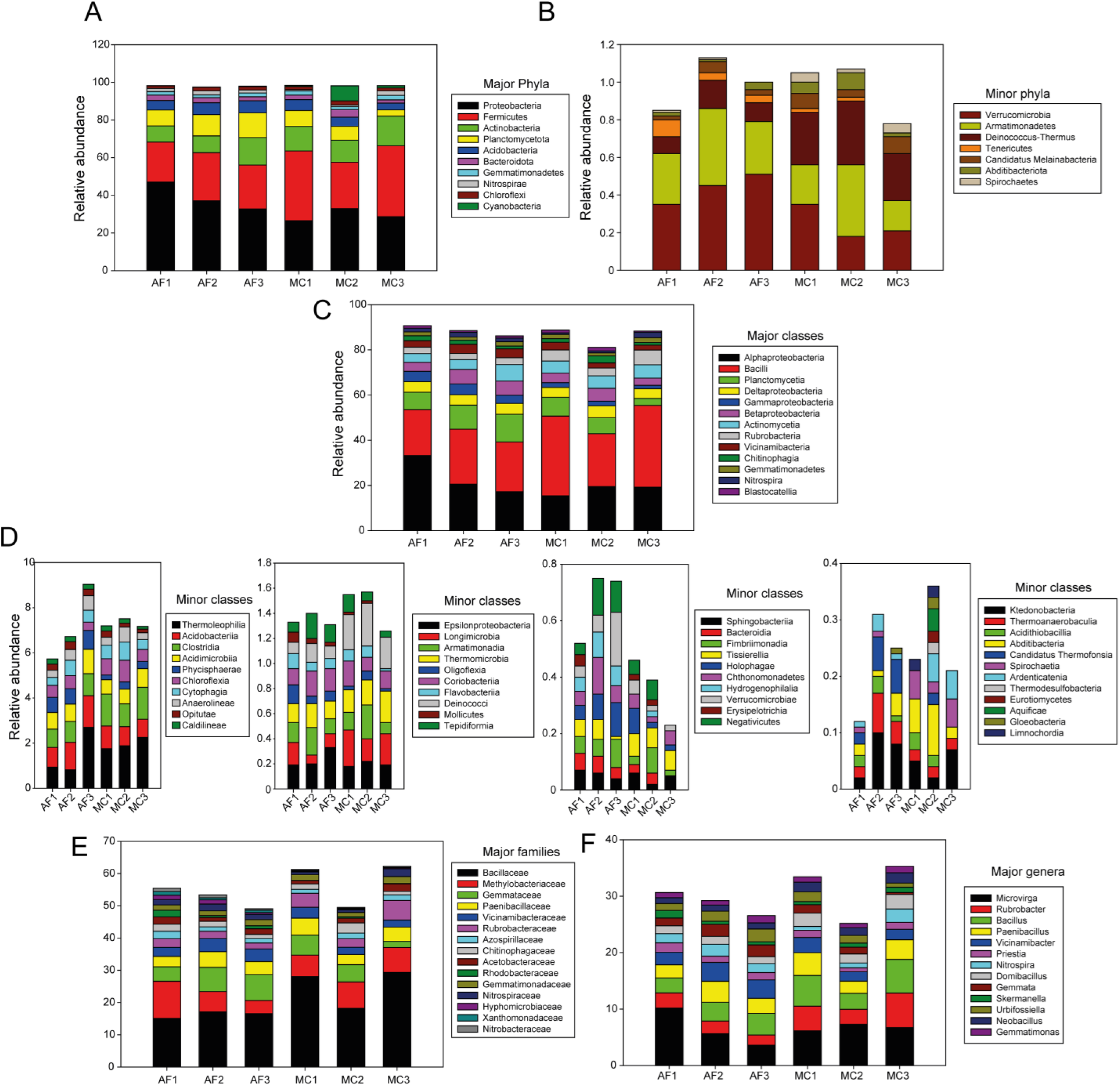
Distribution profile of 16S rRNA gene sequences across different bacterial classifications in rhizosoil from agroforestry and monocropping systems. (A) major phyla, (B) minor phyla, (C) major classes (>1%), (D) minor classes (<1%), (E) major families, and (F) major genera of soil bacteria from the rhizospheres of castor from agroforestry (AF1‒3) and monocropping (MC1‒3) systems. Three replicates from each of AF and MC are shown.

**Table 2.**
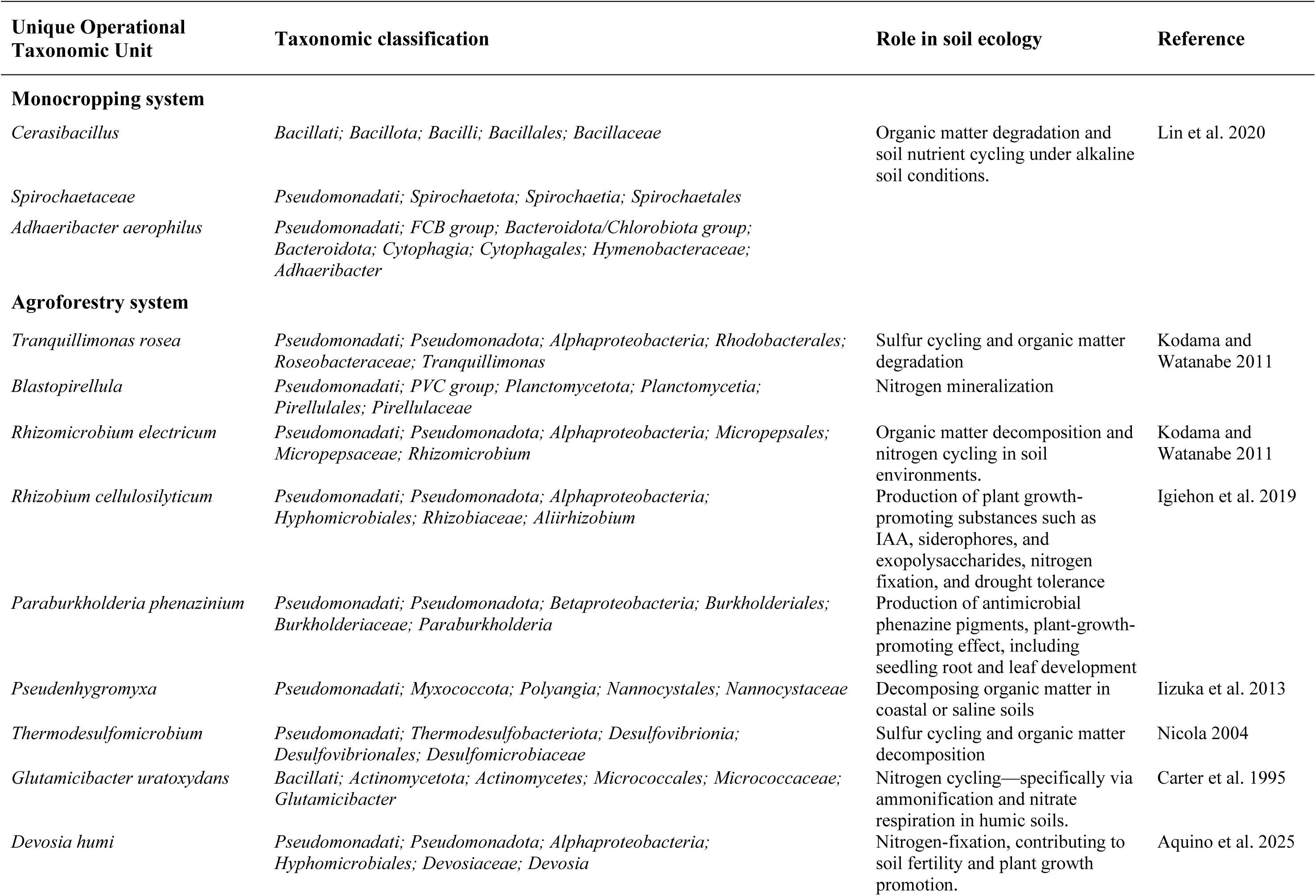
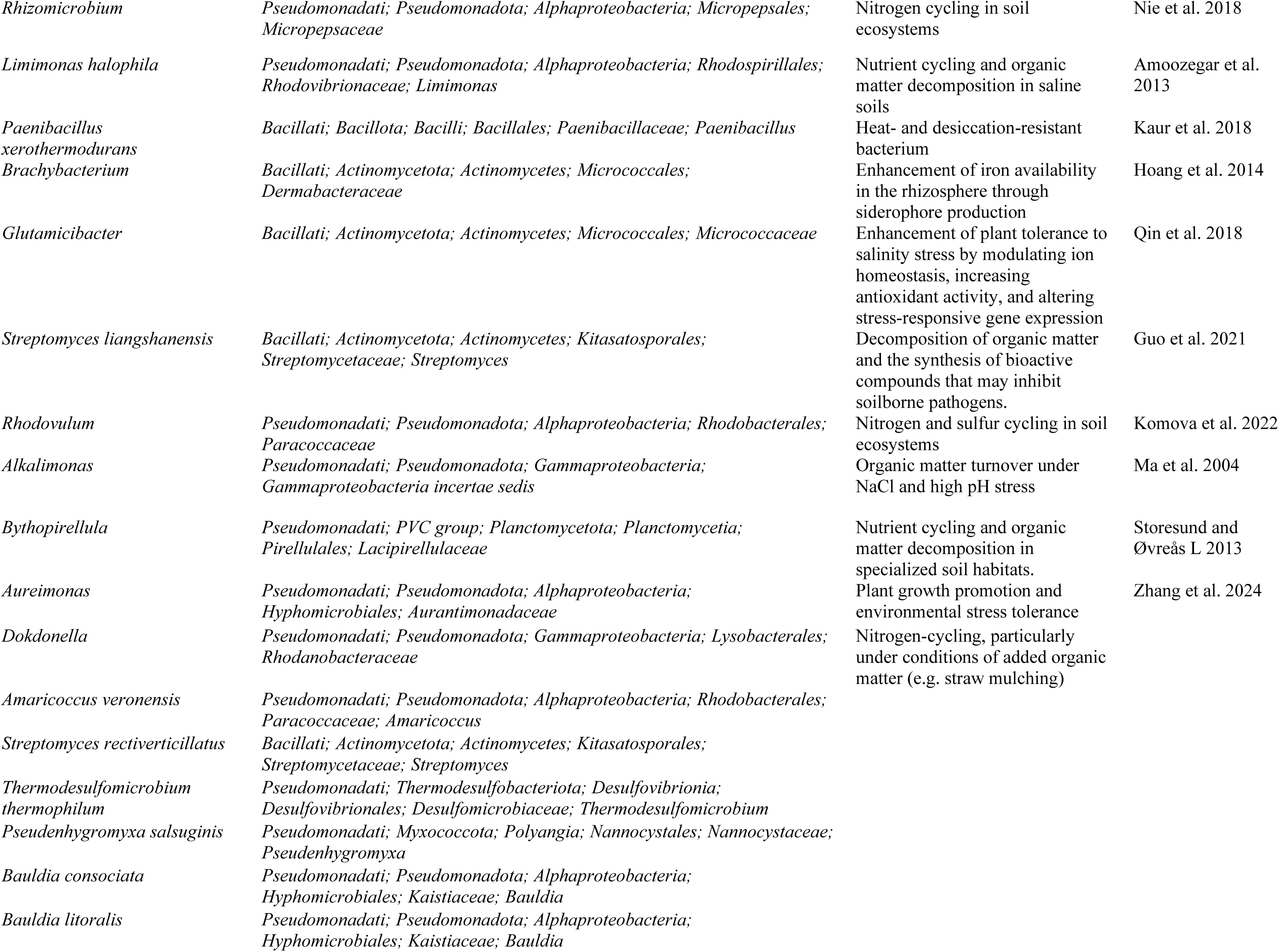
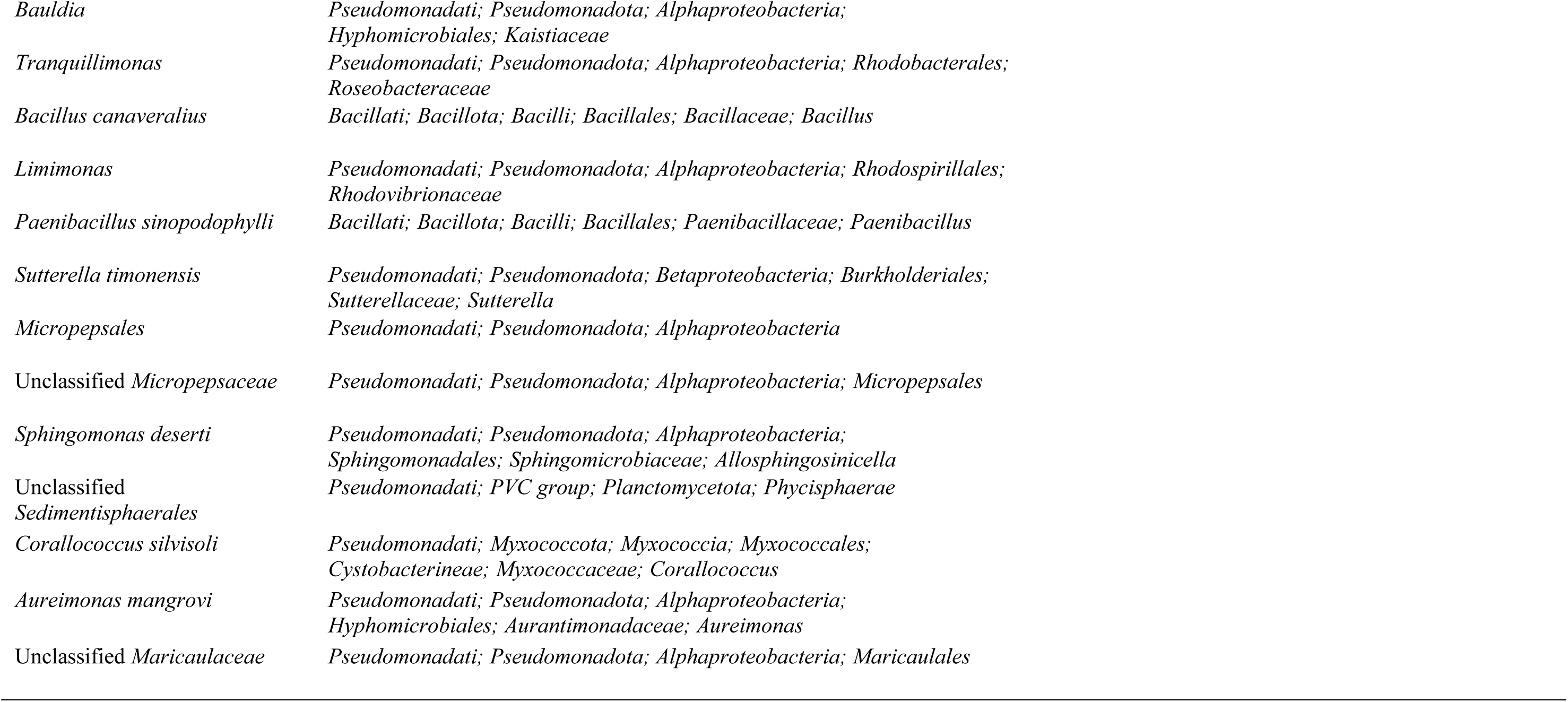
Statistically significant taxa uniquely present in agroforestry and monocropping systems.

### 3.5 Castor rhizosphere and root endosphere contain plant growth-promoting bacteria

We attempted to isolate pure bacterial cultures from the rhizosphere and root endosphere of AF and MC in LB media. On average, AF rhizosphere soil samples yielded a higher (4.6 × 10^6^ CFU g^−1^) number of colonies than MC (3.3 × 10^6^ CFU g^−1^). AF also showed potentially higher root colonization by endophytic bacteria. AF root samples showed a higher colonization rate (2 × 10^6^ CFU g^−1^) than MC (1.3 × 10^6^ CFU g^−1^). The AF system consistently showed higher bacterial abundance than the MC system, indicating that diversified land use supports a more active microbial community. This suggests that AF provides a more favorable environment for root-associated bacteria. Nine distinct endophytic bacteria were isolated from the root endosphere of castor plants grown in AF, alongside seven from the AF rhizosphere. In contrast, six bacteria were recovered from the MC rhizosphere and two from the MC root endosphere. The biochemical characteristics of these pure bacterial isolates were carefully documented (Table 3). The results indicated that most of these bacteria possessed plant growth-promoting (PGP) properties, including phosphate and zinc solubilization, siderophore production, and indole-3-acetic acid (IAA) production. Based on the number of PGP properties, we chose four bacteria, EBAF03, EBAF05, EBAF08, and RCAF01, isolated from AF for further study (Table 3). These four bacteria, EBAF03, EBAF05, EBAF08, and RCAF01, showed significant growth promotion in *A. thaliana* in the soil culture assay, as indicated by rosette diameter and shoot biomass (Fig. 5).

**Fig. 5.**
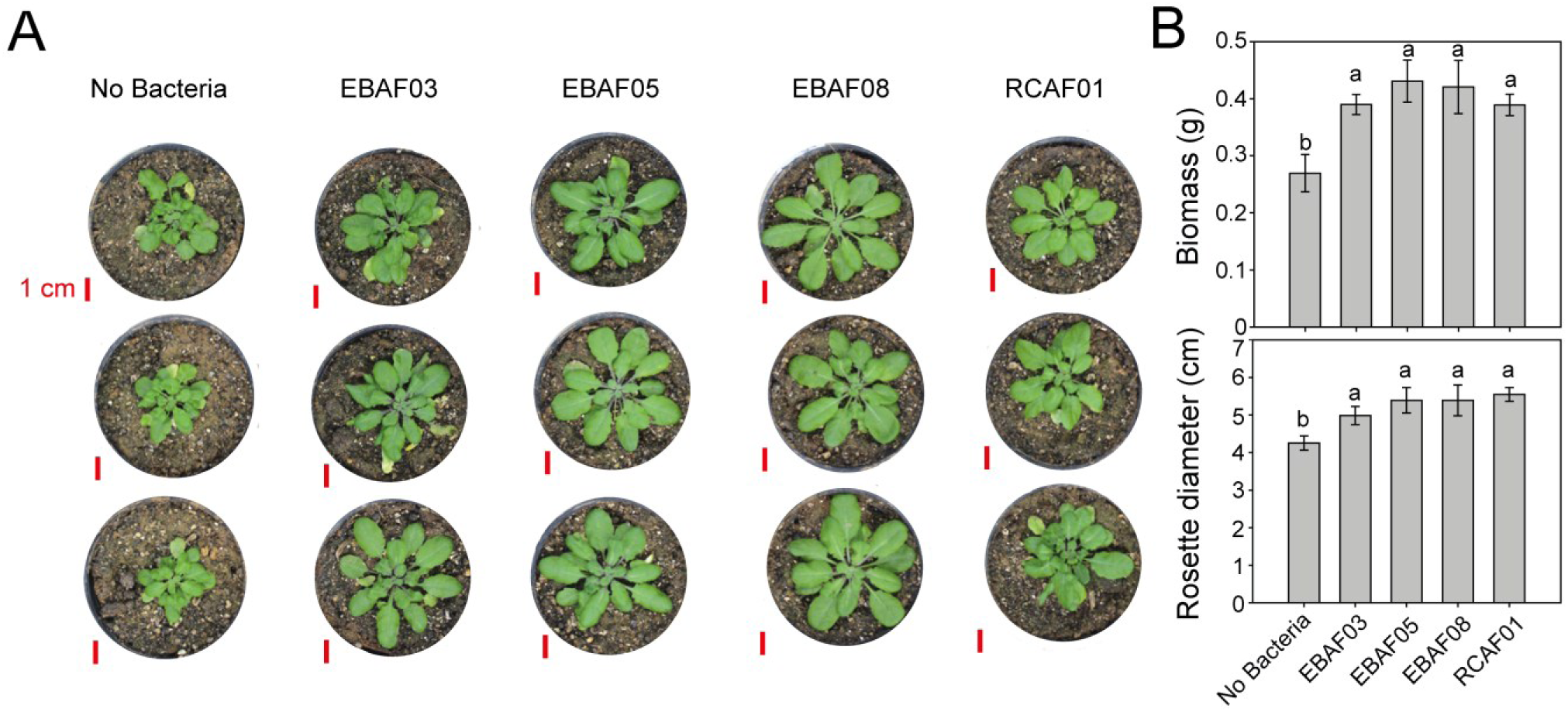
Growth promotion of *Arabidopsis thaliana* by PGPRs from agroforestry-grown castor roots and rhizosphere. **(A)** The effects of plant growth-promoting rhizobacteria (PGPR), EBAF03, EBAF05, and EBAF08 (from agroforestry-grown castor root endosphere) and RCAF01 (from castor rhizosphere) on one-month-old *Arabidopsis thaliana* ecotype Col-0 plants in the soil culture experiment are presented, with three biological replicates per treatment. Red bars represent a scale of 1 cm. **(B)** Average rosette biomass and diameter of six plants in each treatment are presented with standard error after one-month growth in soil. Different lowercase letters above the bars indicate significant differences between samples at *P* < 0.05 (One-way ANOVA followed by Tukey’s HSD-test, *N*=6)

**Fig. 6.**
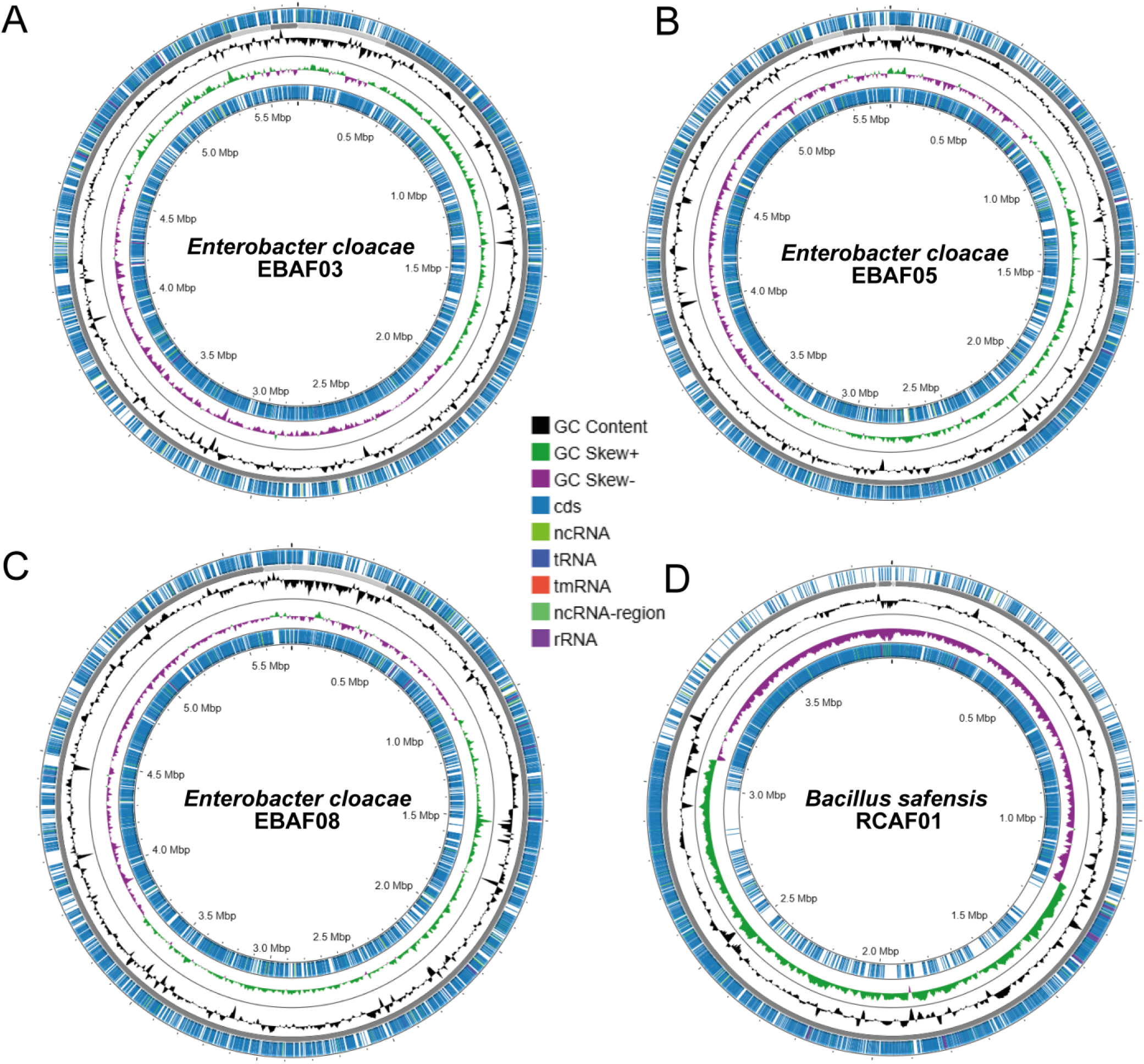
Genome maps of endophytic and rhizospheric plant growth-promoting bacteria from agroforestry-grown castor. *Enterobacter cloacae* EBAF03, EBAF05, and EBAF08, from castor root endosphere and *Bacillus safensis* RCAF0 from castor rhizosphere generated using Proksee. Rings from inner to outer: Bakta-annotated coding sequences (CDSs) on the negative strand (blue), GC skew (+) (green) and GC skew (–) (violet), GC content (orange), other RNAs (per legend), contigs (grey), CDSs on the positive strand (blue).

**Table 3.**
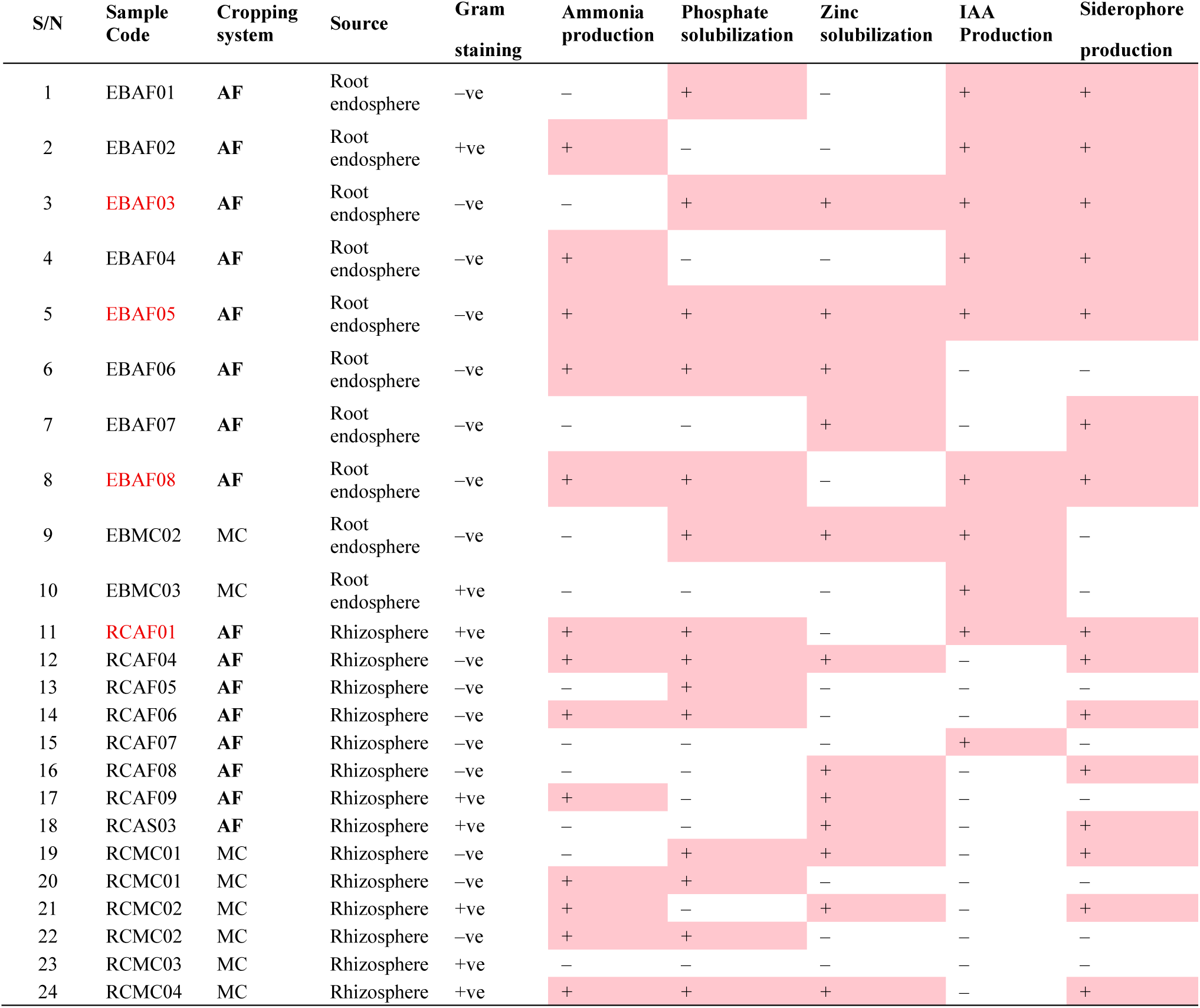
Plant growth-promoting properties of bacteria isolated from castor root endosphere and.

These four bacteria were characterized by whole-genome sequencing. A 16S rRNA gene sequence as well as whole genome-based taxonomic classification analysis revealed that EBAF03, EBAF05, and EBAF08 are different strains of *Enterobacter cloacae*, while RCAF01 was a strain of *Bacillus safensis* (Table S2). *Enterobacter cloacae* EBAF03 harbors a rich repertoire of genes supporting plant growth promotion, as detailed in Table 4. For nutrient assimilation, it encodes enzymes for nitrogen metabolism (e.g., asparagine synthetase, glutamine synthetase, urease subunits), phosphorus acquisition (e.g., glycerol-3-phosphate ABC transporters, polyphosphate kinase, *Phn* phosphonate utilization genes), and sulfur pathways (e.g., sulfite reductase, cysteine synthases). Plant colonization is facilitated by flagellar motility and biosynthesis genes (e.g., *FlhA/B*, *Fli* proteins, *MotA/B*), biofilm formation (e.g., cellulose synthase subunits *BcsB/C*), and chemotaxis (e.g., *CheA/R/Z*, multiple methyl-accepting chemotaxis proteins). Additionally, auxin biosynthesis genes (e.g., tryptophan synthase subunits) and siderophore production/transport genes (e.g., enterobactin cluster: *EntB/F/S*, *FepB/C/D/G*) enhance root growth and iron availability, while fatty acid/oil biosynthesis genes (e.g., acetyl-CoA carboxylase, Fab synthases, acyltransferases) may contribute to lipid-mediated interactions. Similar genes were identified in the two other strains EBAF05 and EBAF08 (Tables S3‒4).

**Table 4.**
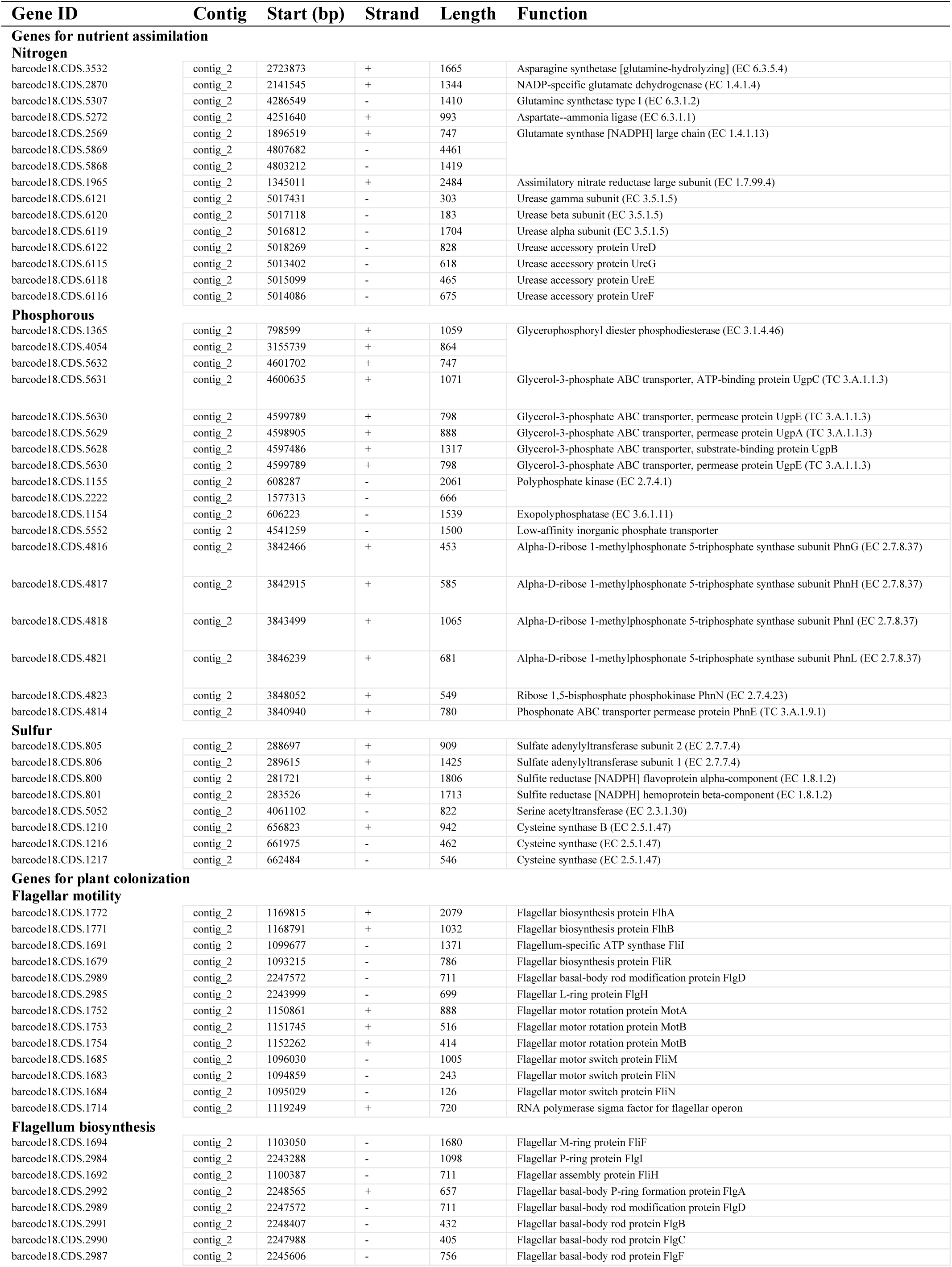
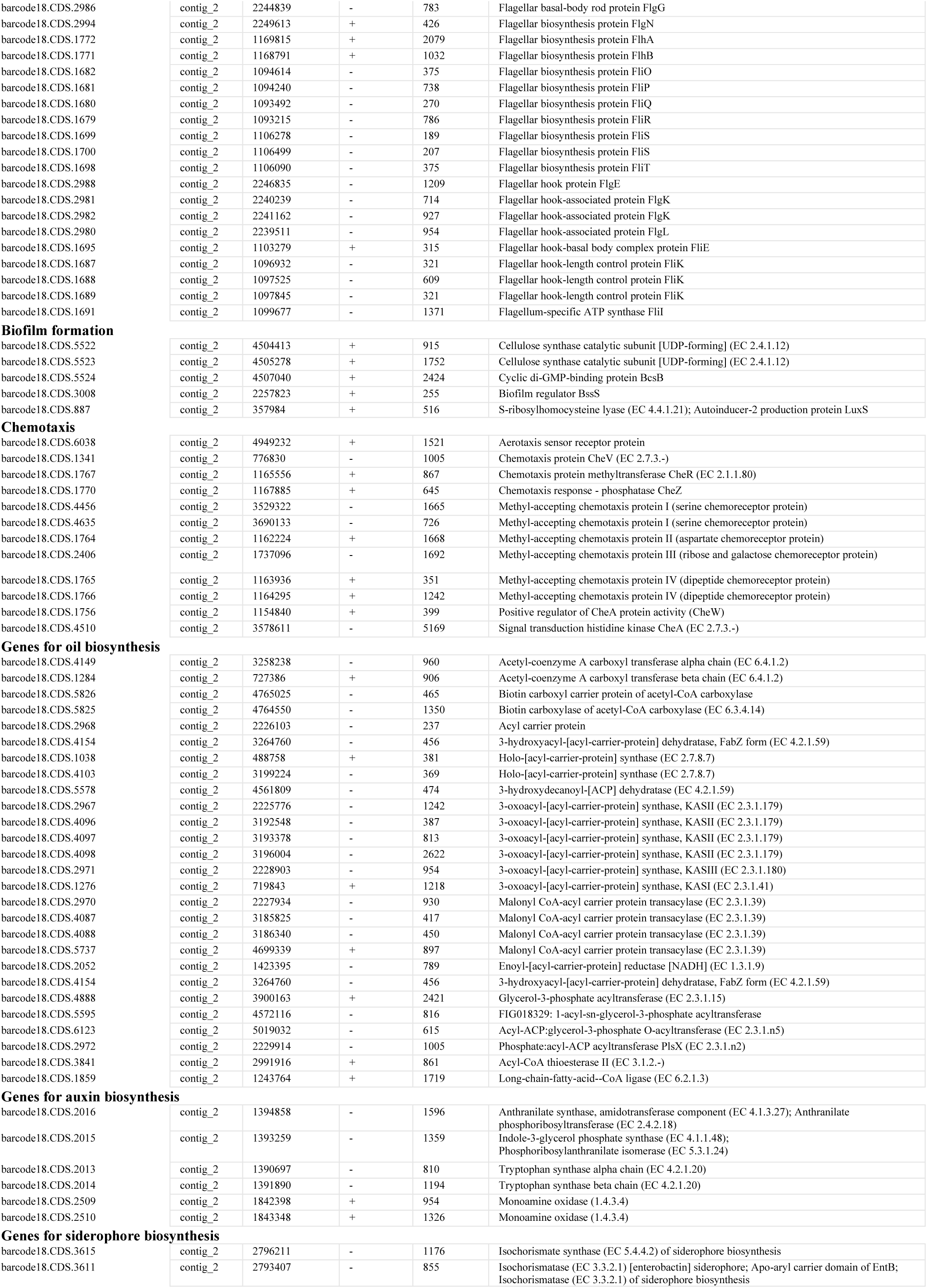
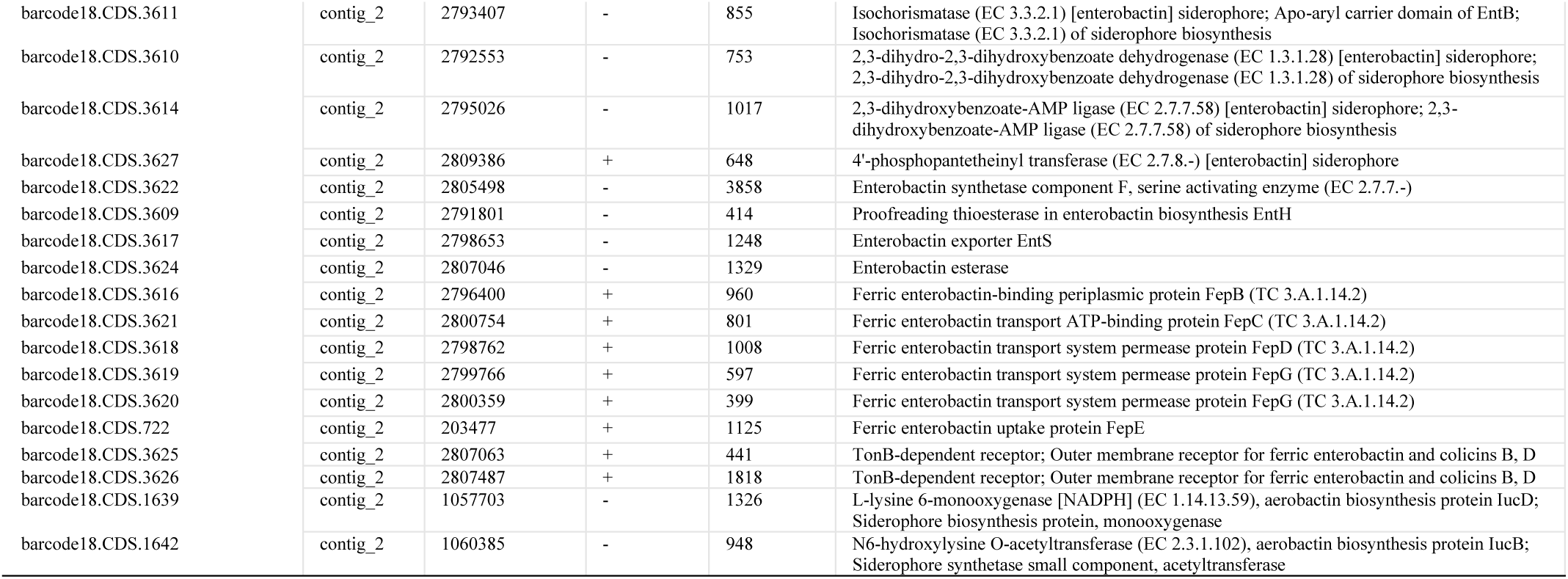
Plant growth promotion and oil biosynthesis genes in *Enterobacter cloacae* EBAF03 genome.

The genome of *Bacillus safensis* RCAF01 encodes diverse genes supporting plant growth promotion through nutrient assimilation, colonization, and phytohormone production (Table 5). Key nutrient assimilation genes include those for nitrogen (e.g., ammonium transporter, glutamine synthetase, multiple nitrate/nitrite reductases), phosphorus (e.g., alkaline phosphatases, *Pst* ABC transporters), and sulfur (e.g., *CysP/T/W/A* sulfate transporters, sulfite reductase). For plant colonization, flagellar motility genes (e.g., *FlhA*, *FliG*, *MotA*/*B*), chemotaxis proteins (e.g., *CheB*, *CheR*), adhesion (fibronectin-binding), and biofilm formation (*TasA*, *SipW*) are prominent. Auxin biosynthesis is supported by tryptophan pathway genes (e.g., anthranilate phosphoribosyltransferase, tryptophan synthases). These features position RCAF01 as an effective PGPR, supporting our observations of plant growth promotion in Arabidopsis (Fig. 5). The genome of *Bacillus safensis* RCAF01 encodes a complete type II fatty acid synthesis (FASII) pathway for oil/lipid production, essential for membrane lipids and biosurfactants that aid rhizosphere colonization. Key genes include acetyl-CoA carboxylase subunits (alpha/beta chains, biotin carboxylase/carrier) for malonyl-CoA formation; multiple malonyl CoA-ACP transacylases; 3-oxoacyl-ACP synthases (*KASII*, *KASIII*); reductases (3-oxoacyl-ACP reductase, enoyl-ACP reductases); dehydratases (*FabZ*); and downstream acyltransferases (*PlsY*) for phospholipid assembly (Table 5). These enable saturated fatty acid chain elongation from C2 to C16/C18, supporting PGPR functions like nutrient emulsification in castor rhizosphere. Since *Bacillus safensis* RCAF01 is a root endophyte of castor, key oil biosynthesis genes in the genome of this bacterium (Table 5) could enhance castor oil quality by influencing host fatty acid metabolism and seed oil accumulation.

**Table 5.**
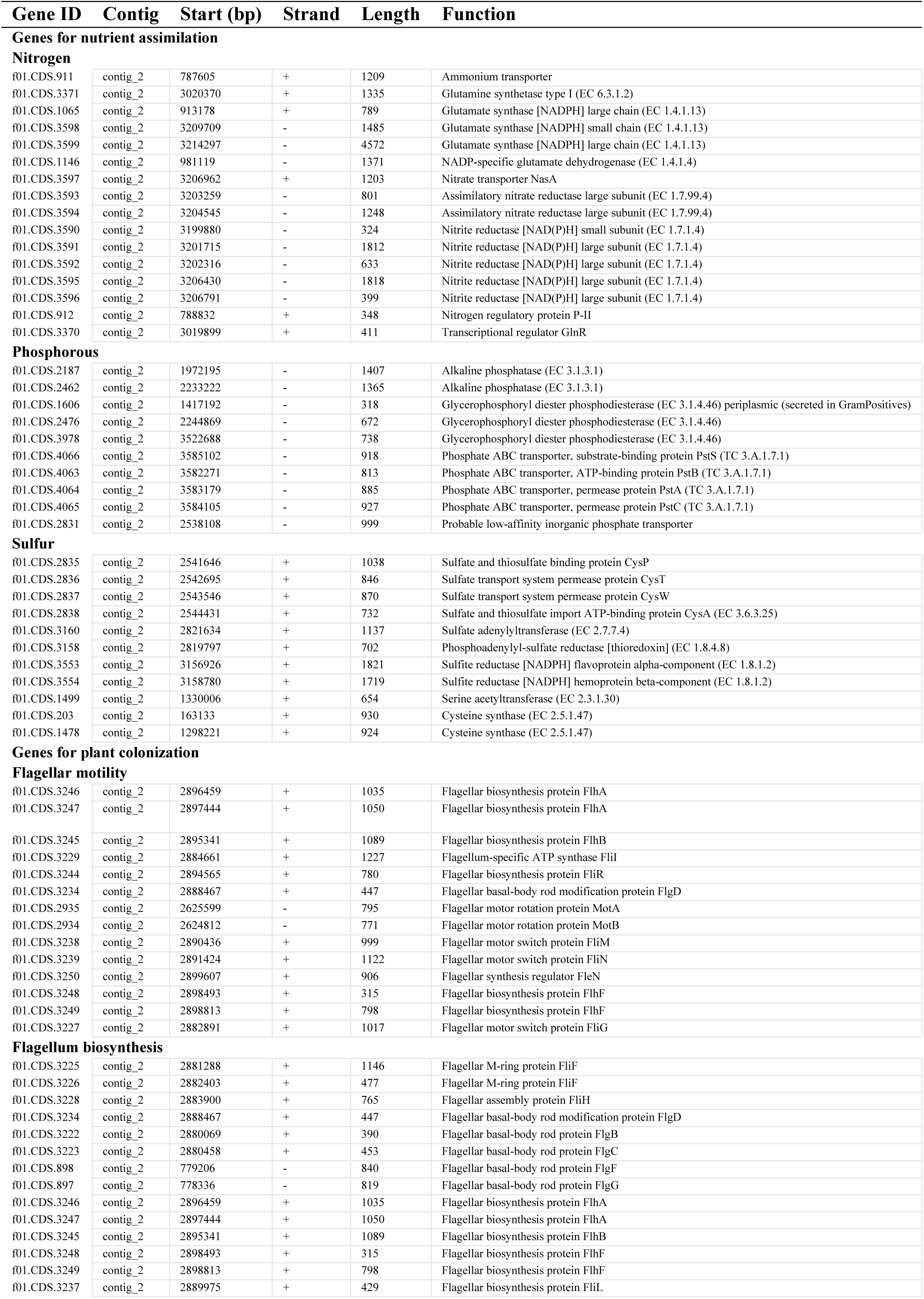
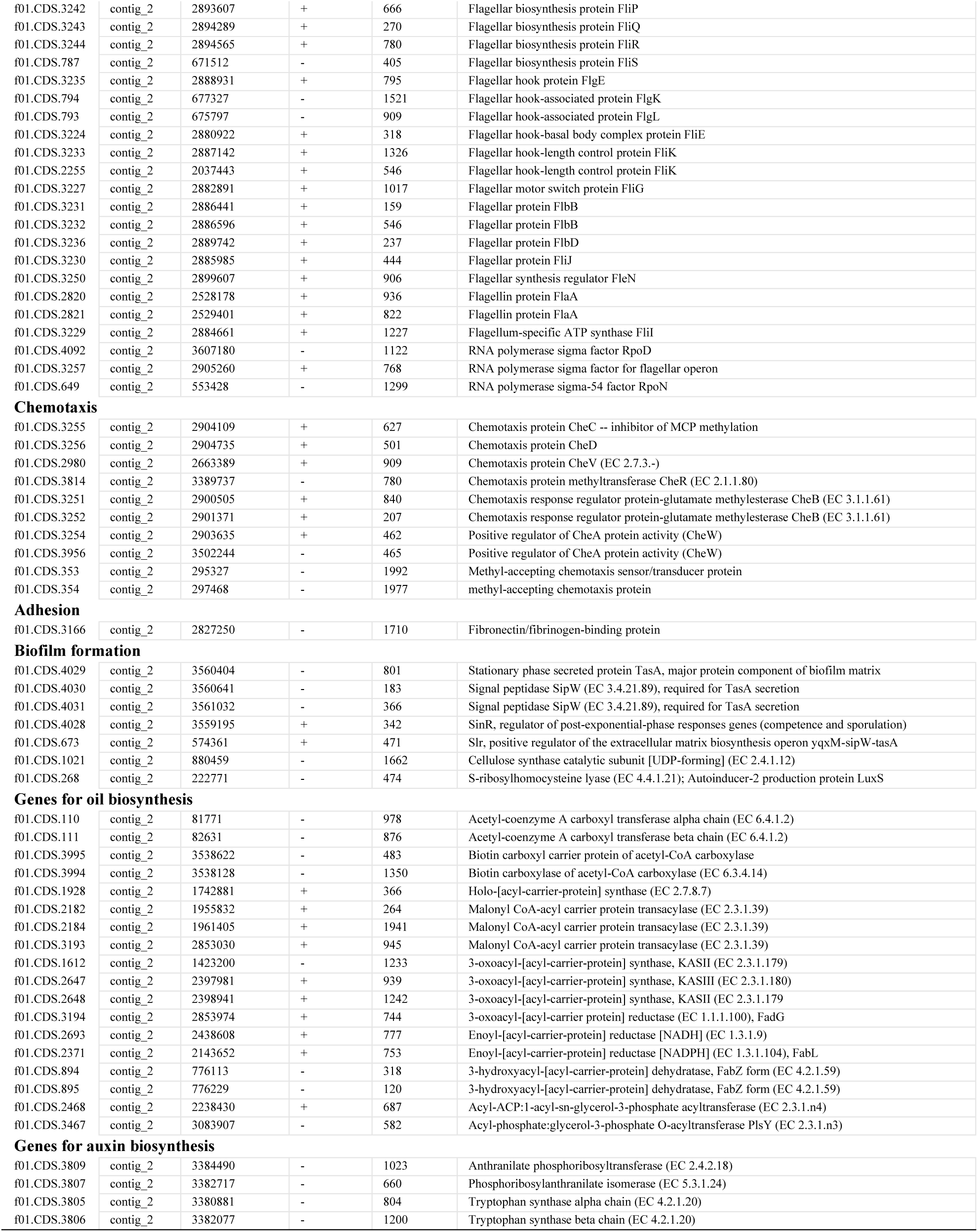
Plant growth promotion and oil biosynthesis genes in *Bacillus safensis* RCAF01 genome.

## 4. Discussion

The present study investigated the antimicrobial efficacy of castor oil derived from *Ricinus communis* seeds cultivated under different systems, AF and MC assessed its concentration-dependent effects against various bacterial strains. These findings are in line with previous research evaluating the antimicrobial properties of castor oil in inhibiting pathogenic microorganisms and its potential applications in medicinal soap production (Abdulrasheed et al. 2015; Aneja et al. 2024).

This study provides a multifaceted insight into the influence of cultivation practices on both soil microbial communities and the bioactive properties of castor oil. The soil microbial analysis revealed a highly diverse bacterial community, predominantly composed of phyla such as *Proteobacteria*, *Firmicutes*, and *Actinobacteria*. Although variations in community composition between MC and AF were observed, these differences were not statistically significant, suggesting a resilient, stable soil microbiome that maintains functional redundancy despite environmental or management differences. At finer taxonomic levels, the higher enrichment of bacteria involved in nitrogen fixation, nutrient cycling, and plant growth promotion, such as *Alkalimonas*, *Aureimonas*, *Blastopirellula*, *Brachybacterium*, *Bythopirellula*, *Devosia humi*, *Dokdonella*, *Glutamicibacter*, *Glutamicibacter uratoxydans*, *Limimonas halophila*, *Paenibacillus xerothermodurans*, *Paraburkholderia phenazinium*, *Pseudenhygromyxa*, *Rhizobium cellulosilyticum*, *Rhizomicrobium*, *Rhizomicrobium electricum*, *Rhodovulum*, *Streptomyces liangshanensis*, *Thermodesulfomicrobium*, and *Tranquillimonas rosea* in the AF rhizospheres (Table 2), underscores the critical role of these microbial consortia in enhancing plant resilience and agroecosystem productivity. The detection of these unique taxa in the AF group suggests the potential to enhance soil health and fertility through cultivation practices such as AF, which may foster beneficial microbial interactions. In previous reports, inoculation with plant growth-promoting bacteria like *Azospirillum brasiliense, Bacillus subtilis*, and *Pseudomonas fluorescens* increased castor yield and oil content (Gato et al. 2024) and aided the remediation of copper and cadmium (Li et al. 2025). Notably, *Alphaproteobacteria*, slightly enriched in the AF rhizosphere, harbor functional genes for fatty acid biosynthesis (*acc*, *fab*), nitrogen fixation (*nif*), reduction of stress-induced ethylene (*acdS*), phosphate solubilization (*gcd*), iron acquisition (*sid*/ *pvd*), and hormone balance (*iaa*) that indirectly enhance castor oil yield and quality by improving nutrient acquisition, stress tolerance, and carbon flux toward fatty acid biosynthesis, particularly ricinoleic acid (Kielak et al. 2016). Genomic analysis of our isolates from AF roots and rhizospheres, viz. *Enterobacter cloacae* and *Bacillus safensis*, revealed the presence of genes related to plant growth promotion (Tables 4, 5, S3, S4). Core fatty acid biosynthesis genes were identified in the genomic analysis of *Enterobacter cloacae* EBAF03, EBAF05, EBAF08 and *Bacillus safensis* RCAF01 (Tables 4 and 5), including *acc* (acetyl-CoA carboxylase complex), *fab* (*KAS*, *ACP*, reductases), and acyltransferases (*PlsX/PlsY/PlsC*) showing a conserved lipid metabolic framework. Similarly, the genes linked with nitrogen assimilation, phosphate uptake, sulfur metabolism, siderophore production and auxin biosynthesis strongly indicate their high plant growth-promoting potential. These traits indirectly boost host fatty acid biosynthetic pathways by improving nutrient acquisition and carbon flux.

On the other hand, a*cetyl-CoA carboxylase* (*RcACC*), *β-ketoacyl-ACP synthase* (*RcKAS*), *diacylglycerol acyltransferase* (*RcDGAT2*), *oleoyl-12-desaturase* (*RcFAD2*), and *Oleoyl-12-Hydroxylase* (*RcFAH12*) are key genes in the castor fatty acid biosynthesis pathway, leading to ricinoleic acid accumulation in seed oil. *RcFAH12* stands out as the signature gene for castor oil quality due to its role in hydroxylating oleate to ricinoleate and to its higher expression in high-oil-producing castor cultivars during seed development (Héctor et al. 2020). A higher induction of *RcDGAT2* in AFS-grown castor regmata compared to MC of the same developmental stage (Fig. 2F) indicates that AFS conditions promote a more specialized oil-filling program: with a higher commitment to ricinoleate-rich triacylglycerol (TAG) formation, potentially improving seed-oil quality (ricinoleate content) relative to MC plots possibly through altered light, water, nutrient dynamics, and rhizosphere/ endosphere microbe interactions creating a resource-partitioning signal that up-regulates this key TAG-assembly enzyme. This shifts the developing seed’s metabolism toward more efficient channeling of ricinoleoyl-CoA into storage oil, enhancing ricinoleate-rich TAG formation (Fig. 2D).

Parallel to these ecological insights, the investigation into the antimicrobial efficacy of castor oil derived from *Ricinus communis* seeds cultivated under different systems revealed complex interactions influenced by cultivation methods. Physical attributes of the seeds, including color, size, and germination rate (Fig. 2A), indicated better overall development in the AF system. Darker seeds with higher vigor and faster germination suggest superior phytochemical content, likely a consequence of improved soil health, organic fertilizers like Jeevamrit, and diversified cropping systems. These factors are known to enhance secondary metabolite production, which directly correlates with the oil’s antimicrobial potential. Susceptibility assays demonstrated that castor oil possesses moderate antimicrobial activity, with zones of inhibition smaller than those of standard antibiotics, but with noteworthy differences across cultivation systems. Interestingly, while agar diffusion tests showed limited inhibition, agar dilution tests indicated that bacterial colony counts were higher in MC systems, implying that environmental factors intrinsic to agroforestry could bolster castor oil’s bioactivity. Furthermore, optical density measurements over 24 hours revealed a paradoxical trend: at higher concentrations, castor oil sometimes promoted bacterial growth, particularly in plant growth-promoting bacteria such as *Bacillus mobilis* and *Pseudomonas fluorescens*. This suggests a dual role for castor oil: an antimicrobial agent at certain concentrations and a nutrient source or growth stimulant at others. The observation that higher concentrations of castor oil stimulated bacterial proliferation, especially within the agroforestry system, raises intriguing questions about the bioactive constituents responsible for these effects. It is plausible that specific phytochemicals or secondary metabolites in the oil, potentially enhanced by organic inputs and diversified plant interactions, modulate microbial activity. This duality underscores the complexity of plant-derived oils in microbial ecosystems: they can suppress harmful pathogens while simultaneously promoting beneficial plant growth-promoting bacteria, thereby contributing to sustainable agriculture. The influence of cultivation practices on the phytochemical profile of castor oil further emphasizes the importance of agroforestry systems. These systems, characterized by diversified cropping and organic inputs, appear to enhance the oil’s bioactive properties, which could be exploited for both medicinal and agricultural applications. Future studies should prioritize the isolation and characterization of key bioactive compounds driving antimicrobial and plant growth-promoting activities, while clarifying their modes of action against a wider array of microbes.

*Bacillus safensis* and *Enterobacter cloacae* isolated from castor rhizosphere and root endosphere in our study are effective PGPRs that enhance productivity in oil-yielding crops like oilseed rape (*Brassica napus*), maize (*Zea mays*), and sunflower (*Helianthus annuus*), as well as other crops such as fenugreek (*Trigonella foenum-graecum*), cucumber (*Cucumis sativus*), and wheat (*Triticum aestivum*) (Gupta et al. 2022; Chebotar et al. 2023; Fan et al. 2025). These bacteria employ multifaceted mechanisms, including indole-3-acetic acid (IAA) production, phosphate and potassium solubilization, nitrogen fixation, exopolysaccharide (EPS) synthesis, and induction of tolerance to abiotic stresses like salinity, drought, and alkalinity. Notably, *E. cloacae* strain PNE2, isolated from vegetable waste, boosts fenugreek seedling vigor through biopolymer degradation and nutrient mobilization, while recent work shows it elevates *Reaumuria soongorica* biomass in desert conditions and enhances saline-alkali resilience in cucumber, cotton, and wheat (Pooja & Patel, 2022; Fan et al. 2025). Endophytic *B. safensis* TS3 colonizes oilseed rape roots, increasing endosphere populations and nutrient uptake under salinity (Chebotar et al. 2023). In oil-contaminated soils, *E. cloacae* subsp. PMEL-79 degrades crude oil while promoting maize root elongation, combining bioremediation with growth benefits, and this is further supported by its biosurfactant production, which leads to oil breakdown (Ejaz et al. 2021). Genomic evidence from our study (Tables 4 and 5) also supports the potential of *Enterobacter cloacae* EBAF03, EBAF05, EBAF08, and *Bacillus safensis* RCAF01 to act as PGPR, as genes involved in nutrient assimilation (N, P, S), siderophore production, and auxin biosynthesis were identified. Such traits increase nutrient availability and root development, thus enhancing carbon allocation and metabolic efficiency in the host plant. These microbially mediated effects could indirectly enhance fatty acid biosynthesis by elevating precursor availability (e.g., acetyl-CoA), thereby contributing to higher oil yield and quality in castor.

In conclusion, this integrated study underscores the interconnectedness between sustainable cultivation practices, soil microbial ecology, and plant secondary metabolite production. Agroforestry not only fosters a resilient and functionally rich soil microbiome but also enhances the bioactive potential of *Ricinus communis* seeds. These findings open avenues for optimizing crop management strategies to produce bioactive compounds with dual benefits, improving soil health and providing natural alternatives for microbial control and plant growth promotion. The dual role of castor oil as both an antimicrobial and growth-promoting agent warrants further exploration, especially in the context of eco-friendly agriculture and medicinal plant-derived products.

## Author Contributions (CRediT Taxonomy)

KS: Conceptualization; Methodology; Supervision; Resources; Writing – original draft. PP: Writing – review & editing. TB: Investigation; Formal analysis; Writing – review & editing. NC: Investigation. AM: Investigation; Formal analysis. RJ: Investigation. RK and PY: Formal analysis. RM: Investigation. PY, NR, and NSC: Investigation; Formal analysis. AS: Methodology; Supervision; Resources; Writing – original draft; Writing – review & editing.

## Raw data

The metagenomic raw data is submitted to NCBI SRA under accession PRJNA1457303, and the whole-genome sequences of the isolated bacteria are submitted under accession PRJNA1456546.

## Acknowledgments and Funding

KS acknowledges funding from Agroforestry Promotion Network Switzerland (AFS-01) in collaboration with Agriculture University Jodhpur. AS acknowledges funding from IIT Jodhpur (I/SEED/ASK/20220015) and SERB (SRG/2022/000169). We also thank the Indian Institute of Oilseeds Research for extending support in GC and NMR analysis of castor oil. Further, the authors acknowledge Maharshi Dayanand University for providing facilities for metagenomic and bacterial whole genome sequencing.

